# Diversity at the *Acinetobacter baumannii* K locus: towards a comprehensive *in silico* database for prediction of capsular polysaccharide types

**DOI:** 10.64898/2026.07.20.739706

**Authors:** Johanna J. Kenyon

**Affiliations:** School of Pharmacy and Medical Sciences, Health Group, Griffith University, Parklands Drive, Gold Coast, Queensland, Australia; Institute for Biomedicine and Glycomics, Griffith University, Parklands Drive, Gold Coast, Queensland, Australia

**Keywords:** *Acinetobacter baumannii*, capsular polysaccharide, K locus, KL, *Kaptive*

## Abstract

*Acinetobacter baumannii* is one of the most critical bacterial pathogens requiring novel approaches for infection control and treatment to curb the spread of highly resistant isolates. Epidemiological surveillance, and the design and application of many non-antibiotic interventions, require rapid and accurate prediction of diverse capsular polysaccharide (CPS) types from whole genome sequence data. The internationally adopted CPS typing system relies on the *in-silico* detection of CPS biosynthesis genes both in and outside the K locus (KL), utilising a reference sequence database that is compatible with the bioinformatics tool, *Kaptive.* In this study, 168 novel loci were added to the database following an extensive survey of publicly available *A. baumannii* genomes and non-redundant sequence entries, bringing the total number of reference sequences to 409 KL and 10 extra-locus genes. All novel loci conformed to the characteristic K locus configuration described previously, with conserved core genes for CPS biosynthesis flanking a central and highly variable region that includes type-specific structural genes. Across the 409 KL, there were 1000 protein clusters. Annotations for 309 novel clusters were curated and validated using a variety of sequence-and structure-based approaches. Most proteins (n=781) were encoded by genes found in ≤4 KL, consistent with extensive structural diversity of CPS types in *A. baumannii* as reported previously. However, gene distribution analysis predicted shared structural features, with several KL predicting the same CPS type or K unit. Validation of the updated database against 46,185 publicly available genomes revealed that 32 K loci were present in 87.1% of sequenced isolates, with an overrepresentation of genomes with KL2, KL3, KL18, KL9 or KL22.

**IMPACT STATEMENT:** Capsular polysaccharide (CPS) typing is embedded in *in-silico* genomic approaches utilised for characterisation and epidemiological surveillance of *Acinetobacter baumannii* isolates. As new information regarding CPS diversity and structures becomes available, continual updates to the integrated typing database are essential to ensure accuracy and endurance as a valuable scientific resource. The rapid expansion of genome data released into public databases, particularly from previously underrepresented geographies and isolation sources in recent years, provided an opportunity to capture further diversity in CPS encoding regions to enhance the utility of the typing database. A total of 168 novel K loci were identified and added to the database, and an analysis of gene distribution across all 409 loci predicted shared CPS features. This work is expected to enhance KL typing and prediction of CPS structural features for studies involving diagnostics, surveillance and therapeutics applications.

**DATA AVAILABILITY:** All genome sequences used in this study are publicly available with details and accession numbers listed in Supplementary Table 1. The updated database with 409 KL and 10 extra-locus reference sequences is freely available at https://github.com/johannajkenyon/Abaumannii_surface_polysaccharide_loci.

## INTRODUCTION

While globally recognised as one of the leading causes of multidrug-resistant bacterial infections worldwide (1, 2), *Acinetobacter baumannii* has been increasingly identified across diverse ecological niches, highlighting the importance of a One Health approach to track and control isolates resistant to clinically available antibiotics (3–5). Accurate diagnostics, targeted surveillance and outbreak investigations often rely on prediction of the type of structurally variable polysaccharide capsule (CPS) on the cell surface. Many innovative therapeutic approaches, including vaccines (6), monoclonal antibodies (7–9), bacteriophage and their products (e.g. 10-16), also have targeted recognition for specific CPS types. Hence, identification of the many forms of CPS produced by *A. baumannii* is critical across several areas. As rapid serological methods are not available for this species, CPS types are deduced via an *in-silico* approach that is based on the detection of CPS biosynthesis genes in whole genome sequence (WGS) data (17, 18).

In *A. baumannii*, genes relevant to CPS typing are primarily found at the K locus (KL), which is located between conserved *fkpA* and *lldP* genes on the chromosome (Fig. 1a; (19)). To date, sequences found at this location have adopted a general configuration of three defined regions (17–19). Region 1 harbours conserved core genes for CPS export machinery (*wza-wzb-wzc*), which are divergently transcribed from the remainder of the locus. Region 3 includes conserved core genes for synthesis of common sugar precursors (*galU, ugd, gpi*, *pgm*) and may also include a *gne1* gene that is needed to generate nucleotide-linked D-galactose (D-Gal) or N-acetyl-D-galactosamine (D-GalNAc) precursors (19, 20). These regions are either side of Region 2 that generally includes type-specific structural genes, such as those for the synthesis of complex nucleotide-linked sugars, initiating transfer, glycosyltransfer, modification, and CPS processing via the Wzx/Wzy-dependent pathway (19). In a small number of cases, the *wzy* polymerase gene is missing from Region 2, and either sits within the locus (i.e. between *fkpA* and *lldP*) to the left of Region 1 (17, 19, 21), or is found in one of two known genomic islands, referred to as GI1 (22) and GI2 (23).

**Figure 1.**
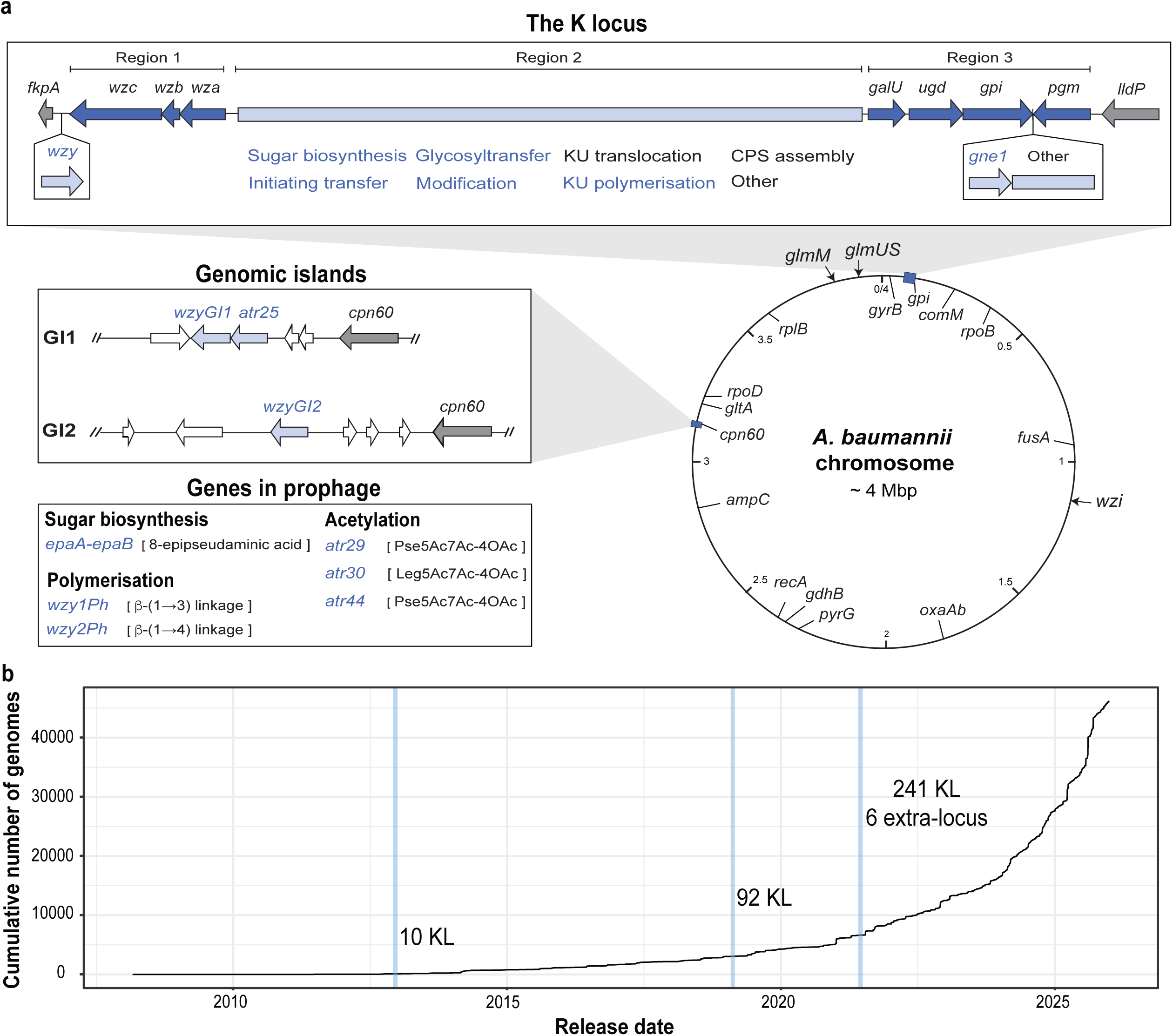
*In-silico* typing of *A. baumannii* CPS. **(a)** Genes relevant to CPS typing in *A. baumannii*. Representation of a chromosome map is bottom right, indicating positions of the K locus, genomic islands and other genes involved in CPS biogenesis outside the circle. Other diagnostic markers, including *oxaAB* and Institut Pasteur and Oxford MLST alleles, are shown inside circle. General features of the K locus are shown above, and known extra-locus genes are shown in the boxes to the left. Conserved genes are dark blue and variable genes light blue. Names in blue font are those involved in determination of CPS type. **(b)** Number of genomes released into NCBI databases per year (March 2007 to 1^st^ Jan 2026). Blue vertical bars indicate key studies reporting characterisation of K loci in available *A. baumannii* genomes conducted in 2013 (19), 2019 (18) and 2021 (17).

To catalogue distinct genetic arrangements, consecutive KL numbers are assigned to sequences based on discernible differences in gene presence in one or more of the variable portions of the K locus (light blue regions in Fig. 1a; allelic differences in conserved core genes do not impact KL assignments (17)). Use of this definitive nomenclature scheme relies on the classification of genes using a translated protein identity threshold of 85lJ% and the assignment of gene names that in most cases reflect the function of the encoded products (i.e. *itr* for the initiating transferases, *gtr* for glycosyltransferases, etc.). This scheme is in active use for KL typing and CPS gene nomenclature in *A. baumannii*, and a complete list of annotations described to date is available in Cahill *et al*. (17).

Where structural data is available, the system for KL numbering extends to the designation of ‘K’ numbers for CPS types. For example, a CPS type will be named K2 if it is produced by an isolate that carries the KL2 locus (24). However, there are some exceptions to this general rule. First, Region 3 may include additional genes between *gpi* and *pgm* that have no established role in CPS biosynthesis but warrant the assignment of a new KL number. Hence, some pairs of KL that differ in this region have been shown to produce the same CPS though they carry different KL, e.g. KL3 and KL22 both produce K3 CPS (25, 26). In addition, the interruption of genes due to frameshifts (26) or insertion sequences (25) may direct structural changes, and in such cases, a numbered ‘v’ suffix is assigned to indicate that the CPS is a variant of a known type (e.g. K3-v1). Finally, as CPS can be changed by ‘extra-locus’ genes found in prophage (27–31) (Fig. 1a), a suffix is also assigned to the name (e.g. K58-8ePse) to indicate the structural change.

The number of catalogued KL has continued to rise since the first detailed characterisation of the K locus in 2013 ((19), Fig. 1b). In 2019, a total of 92 KL were collated into a database compatible with the *in-silico* typing tool, *Kaptive* (18), and this database was updated in 2021 to include 149 further loci (241 KL total) that were identified in genome assemblies available at that time (Fig. 1b; 17). Six extra-locus genes were also added to this database, and an additional special logic file was released to enable the prediction of K type from the specific combination of KL and extra-locus genes detected (17). Since the development and subsequent update of the database, four additional extra-locus genes have been identified (27, 29–31). There has also been a substantial increase in the number of publicly available genome sequences (Fig. 1b), including those from previously underrepresented geographical regions and isolation sources (Supplementary Fig. 1), which presents an opportunity to uncover further CPS biosynthesis genes and increase the utility of CPS typing.

Here, an update to the *A. baumannii* CPS typing database and associated special logic is reported to incorporate novel sequences found in 46,185 publicly available genome sequences surveyed in this study and in NCBI non-redundant entries. The update was applied to assess the frequency of KL amongst sequenced isolates, and the distribution of genes at the K locus is also reassessed.

## METHODS

### Collation and quality filtering of *A. baumannii* genome assemblies

*A. baumannii* genome assemblies (n= 47,214 available under taxid: 470, 1^st^ January 2026) were downloaded from the NCBI using *NCBI datasets v 18.5.0* (32). Associated assembly statistics and metadata (i.e. country of isolation, collection date, isolation source) were either downloaded directly from Biosample records or using metatools_ncbi (https://github.com/farhat-lab/metatools_ncbi). Genome assemblies were retained for analysis if they met the following quality and inclusion metrics: a) the total genome size is between >3 Mb and <4.7; b) N50 (shortest contig length that together with longer contigs, collectively represents 50% of the total genome length) is >20 kbp; c) %GC content is between 38.5% and 41.5%; d) the total number of contigs is <1100; and e) not identified as a sequenced copy of a laboratory reference isolate. Remaining assemblies were screened for the presence of the intrinsic *oxaAb* gene (also referred to as *bla_OXA51-like_*) using ABRicate (https://github.com/tseemann/abricate) to confirm the *Acinetobacter baumannii* taxon.

Following filtering, a total of 1,029 assemblies were removed, leaving 46,185 assemblies for downstream analyses. Details of the genome assemblies, including all metadata and sequence statistics, used in this study are available in Supplementary Table S1.

### Identification of novel K locus reference sequences

*Kaptive v 3.1.0* (klebgenomics.github.io/Kaptive/; (33)) was used to screen the filtered pool of *A. baumannii* genome assemblies (n=46,185). This was performed using the reference sequence database, that includes 241 KL and 10 extra-locus genes relevant to CPS typing (17). The following scoring and confidence parameters were implemented to assist with detecting novel loci: a) `--min-cov 85` defining 85% minimum gene coverage to identify best match sequences; b) `--gene-threshold 85` defining the minimum translated protein identity threshold as 85%, consistent with the criteria used for the *A. baumannii* CPS nomenclature system (19); c) `--max-other-genes 0` defining the locus as ‘Typeable’ if 0 other genes are detected; d) `--percent-expected 100` defining the locus as ‘Typeable’ if 100% of ‘expected’ genes are present.

The results of this search were filtered to remove assignments with fragmented sequences or those with a contiguous match to a KL reference sequence with no additional genes, missing genes, genes below the translated protein identity threshold of 85%, and with a length discrepancy less than +/-500 bp. The K locus was manually inspected in the remaining assemblies to identify those with a potentially unique combination of genes. These sequences were then iteratively compared against established related KL reference sequences using blastn or *pygenomeviz v 1.6.1* (https://github.com/moshi4/pyGenomeViz) with the CLI BLAST workflow implementing an 85% protein identity threshold. Additional novel KL sequences available in NCBI non-redundant entries were also collected for addition to the database. A representative reference for each confirmed novel sequence was selected and assigned a KL number (i.e. KL242-KL409) in the *A. baumannii* KL nomenclature scheme (17–19). In cases where a reference sequence was interrupted by an insertion sequence (IS) and an alternative IS-free sequence was not available, the IS was manually removed from the locus sequence as described previously (18). Details of all reference sequences included in the database are available in Supplementary Table S2.

### Annotation of novel K locus reference sequences

*PPanGGOLiN v 2.2.6* (https://github.com/labgem/ppanggolin/;(34)) was used to generate a gene presence/absence matrix for the complete set of 409 KL reference sequences. This was performed using a translated protein identity threshold of 85% with the `--mode 2` MMSeqs2 algorithm (https://github.com/soedinglab/MMseqs2/wiki#clustering-modes). As annotation of *A. baumannii* CPS genes has traditionally been performed via alignment with characterised references rather than sequence clustering (19), further manual curation was conducted to verify separation of coding sequences as novel groupings. Representative protein sequences for curated clusters were then assessed for protein domains by screening against the Pfam-A database using *hmmscan* via the HMMER *v 3.4* package (35)) with a bitscore threshold of 25. Superfamilies were identified using *Superfamily 2* (36), and Carbohydrate-Active EnZyme (CAZy) domains identified using dbCAN3 (37). Transmembrane segments were predicted using TMHMM2.0 (https://services.healthtech.dtu.dk/services/TMHMM-2.0/), and families for Wzy polymerases confirmed using Hidden Markov Models (HMMs) specific for Wzy, as recently described (38). The top hit was recorded for each protein, with collated results available in Supplementary Table S3.

All assignments were supported by additional primary sequence and tertiary structure alignments. Further clustering with lower amino acid sequence identity thresholds (30%, 40%, 50%, 60%, 70% and 80%) was performed using *PPanGGOLiN v 2.2.6* as described above. BLASTp was used to identify homologues of known or predicted function in public databases. Tertiary structures (monomers) were modelled using *AlphaFold v3* (39), and pTM template modelling scores for model_0 of each protein are listed in Supplementary Table S3. Structural alignments were performed using *Foldseek v 10.941cd33* (40) against a database of available PDB entries, or using the Foldseek Search server (https://search.foldseek.com/) for proteins of unknown function. Annotations for novel sequences were assigned in accordance with the *A. baumannii* CPS nomenclature system (17–19).

### Frequency and distribution of CPS genes across K loci

The final curated matrix was used to plot the distribution of genes across all K loci, and cluster KL reference sequences based on Region 2 gene presence/absence. The optimal algorithm for hierarchal clustering using agglomerative methods (single, complete, average, ward.D2) was calculated using the *cluster* and *factoextra* packages as described previously (41). This was determined to be ward.D2, which was implemented for all further analyses. Plots were constructed using *ggplot2* (42) and *pheatmap* (https://github.com/raivokolde/pheatmap) packages in RStudio. Visual representations of all K loci were drawn using *pygenomeviz v 1.6.1* (https://github.com/moshi4/pyGenomeViz) and annotated in Adobe Illustrator.

### Validation of an updated *A. baumannii* KL reference sequence database

Annotated reference sequences for new loci (KL242-KL409) were incorporated into the *A. baumannii* primary CPS reference database, providing a final total of 409 KL and 10 extra-locus genes. The database file was also updated to include K types and citations for associated structural data published since the last update. In cases where a structure for a specific KL has not yet been determined, the reported ‘K type’ specified in the *Kaptive* output remains defined as “unknown”, as described previously (17). Both the previous database (assigned database *v 3.2.2*) and updated database (assigned database *v 3.3.0*) are available at https://github.com/johannajkenyon/Abaumannii_surface_polysaccharide_loci. To verify the performance of the updated database, *Kaptive v 3.1.0* was rerun on the pool of 46,185 assemblies using the parameters described above. All plots were constructed using *ggplot2* (42) in RStudio. Where extra-locus genes were found in prophage, the region surrounding the gene(s) was characterised using PHASTEST (43) and via alignment with the A320 reference genome (NCBI accession number CP032055.1), as described previously (29, 31). At the conclusion of this study, the special logic file was converted to TOML format specifying the phenotype logic required for *Kaptive* typing and is also available at https://github.com/johannajkenyon/Abaumannii_surface_polysaccharide_loci.

## RESULTS

### Identification of novel CPS biosynthesis genes in *A. baumannii* genomes

Following filtering using specified quality metrics (see methods), 46,185 *A. baumannii* genome assemblies were screened against 241 KL and 10 extra-locus gene references using *Kaptive v 3.1.0.* A total of 35,451 (76.8%) assemblies could be assigned a contiguous match to a known sequence (Supplementary Table 4). These included several KL that were interrupted by 1 or more insertion sequences (IS, n=403 genomes) or an insertion (>4 kbp) that resembled a transposon carrying genes predicting proteins unrelated to CPS biosynthesis (n=28). A further 9,883 (21.4%) assemblies had a discontinuous match, indicating fragmentation of the K locus across multiple contigs. A sequence at the K locus could not be verified for an additional 66 genomes that had evidence of low sequence or assembly quality, and seven genomes that had no detectable K locus. These genomes were set aside and not further evaluated. Amongst the remaining 778 assemblies, 163 novel K loci were identified. Together with an additional 5 novel loci available in NCBI non-redundant entries, a total of 168 novel KL were collated, bringing the total number of unique KL found in *A. baumannii* to 409. Consistent with previous observations (17), all sequences ranged in length from 18.5 to 36.7 kbp (median=24.8 kbp) and were comprised of 16-32 open reading frames (median=21).

### Annotation of CPS biosynthesis genes

Sequences of encoded gene products (n=8817) from 409 K loci were clustered into homologous groups using the 85% translated protein identity threshold prescribed for annotation of CPS biosynthesis genes in *A. baumannii* (19). Curation of the dataset together with pre-existing annotations revealed a total repertoire of 1000 protein clusters (Supplementary Table 3), including 309 novel clusters. To predict their roles in CPS biosynthesis, a variety of tools were used to infer function from sequence relationships. Protein sequences for the 10 documented extra-locus genes were also included in the analysis to assist with functional prediction of novel clusters, providing a total repertoire of 1010 protein clusters. Representative sequences for most clusters could be characterised via detection of one or more domains for protein families (Pfam-A), superfamilies or CAZy families (Fig. 2a). However, detection of ≥6 transmembrane segments (TMS) was often necessary to identify integral membrane proteins, Wzx and Wzy, that did not include a detectable domain, and these were then differentiated from each other using HMMs specific for Wzy (38). Only 13 sequences had no identifiable domains or included ≥6 TMS (Fig. 2a).

**Figure 2.**
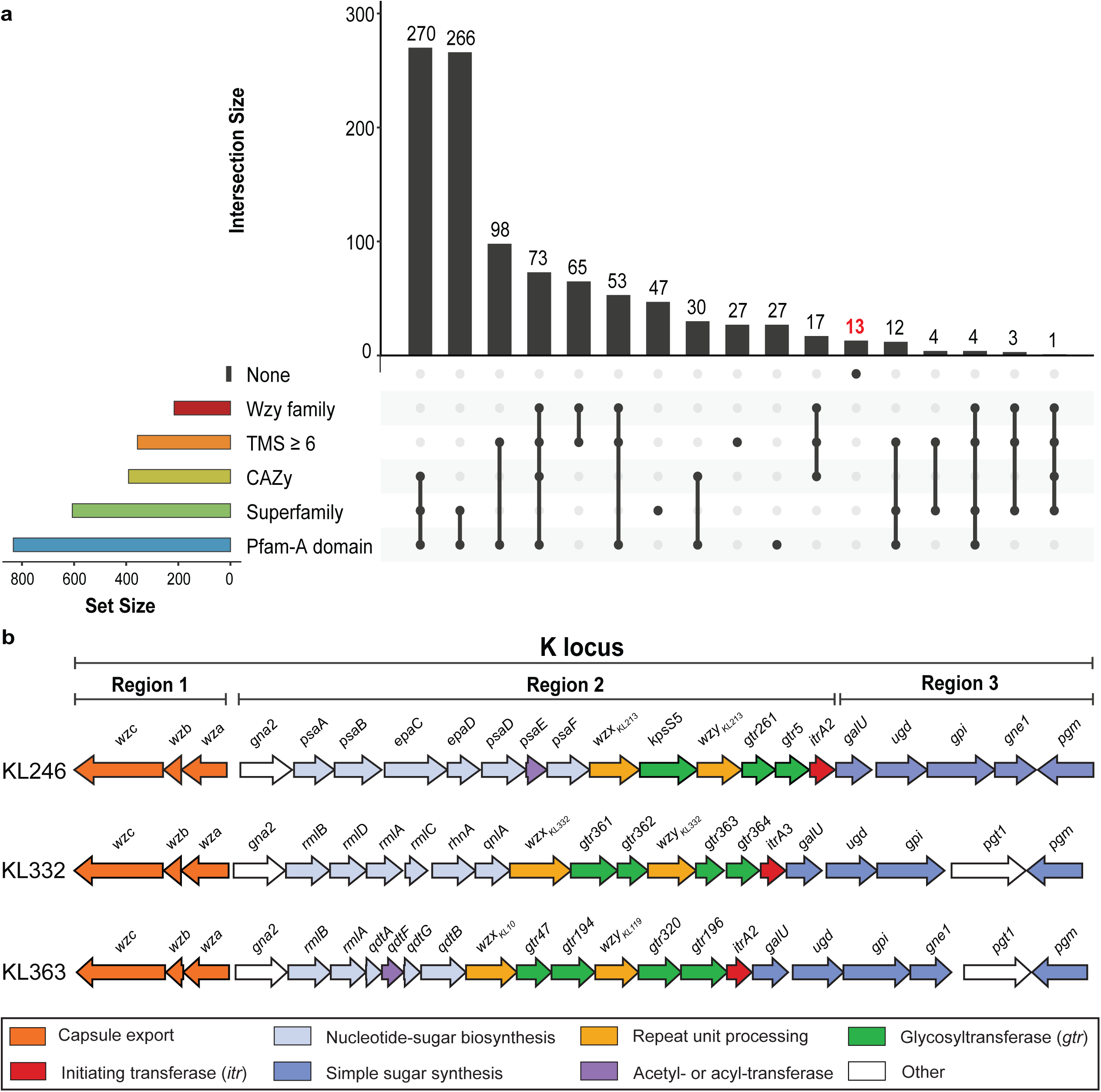
Annotation of CPS biosynthesis genes in *A. baumannii* genomes. **(a)** Intersection plot showing the utility of sequence similarity methods for CPS gene annotation. Number of sequences with hits per tool is shown by horizontal bars to the left. Number with hits for multiple tools is shown by vertical bars above with each intersection shown as connected black circles below. **(b)** K loci with gene modules predicting the synthesis of a novel sugar. Genes are represented as arrows in direction of transcription. Colours are functional categories of gene products and scheme is below.

Additional sequence clustering at lower thresholds (30-80% aa) and searches for homologous products of known or predicted function from other species supported the assignments and allowed further sequences to be functionally predicted. However, to verify predictions based on sequence properties and relationships, tertiary structures for single representatives of all 1010 clusters were also modelled using Alphafold3 and screened against a database of publicly available Protein Data Bank (PDB) entries. Structural relationships with TM-scores >0.5 could be inferred for 706 clusters, with most of the remaining clusters (n=213 of 304) representing predicted Wzy polymerases (Supplementary Table 3). A protein, annotated as GnlB, previously predicted to be involved in synthesis of L-GalNAc (17), was found to be structurally related to a heparinase and belonged to the Chondroitin AC/alginate lyase superfamily, suggesting a different functional role. For products of unknown function, one sequence (previously annotated as Orf_KL68_) was found to be structurally similar to a catalogued glycosyltransferase and was reassigned accordingly. Taken together, a total of 16 clusters could not be assigned roles in the CPS biosynthesis pathway and were annotated as hypothetical proteins. The associated genes were in either Region 2 (n=11) or Region 3 (n=5) between *gpi* and *pgm*. Further searches revealed that 10 of the 16 clusters were structurally related to hydrolase enzymes. However, as their function(s) in relation to CPS biosynthesis are unclear, their annotations as unknown proteins (i.e. Orf_KL#_) were retained.

Collectively, the assignments inferred from sequence- and/or structure-based approaches allowed all proteins to be classified into one of ten functional categories, namely: (i) sugar biosynthesis; (ii) initiating transfer; (iii) glycosyltransfer; (iv) Wzx translocation; (v) Wzy polymerisation; (vi) pyruvyltransfer; (vii) acetyl-/acyl-transfer; (viii) D-alanine transfer; (ix) CPS export machinery; and (x) other. Novel gene sequences included ones for glycosyltransferases (designated *gtr274-gtr426;* n=153), acetyl-/acyl-transferases (*atr45-atr65*; n=21), pyruvyltransferases (*ptr9-10*; n=2), Wzx translocases (*wzx_KL#_*; n=35), and Wzy polymerases (*wzy_KL#_*; n=77). Three novel genes, named *mnaA5-mnaA7*, were found in seven different KLs and are predicted to encode enzymes for the synthesis of UDP-N-acetyl-D-mannosamine (UDP-D-ManNAc) or a derivative thereof. There were also several new homology groups for Wza, Ugd, Gpi, FnlA, FnlB, RmlC and RmlD enzymes not previously reported (17) that fell below the 85% protein identity threshold, which may indicate the acquisition of these genes from a source other than *A. baumannii*.

Amongst the 168 novel loci, three KL (Fig. 2b) included novel gene(s) for the synthesis of a sugar not yet identified in an experimentally verified *A. baumannii* CPS structure. The first of these, KL264, harboured a gene module that included known *psa* genes for the synthesis of CMP-5,7-di-N-acetylpseudaminic acid (CMP-Pse5Ac7Ac). However, *psaC* had been replaced by two genes, designated *epaC* and *epaD*. These genes code for products that respectively share 42.5% and 56.7% to EpaA and EpaB for conversion of Pse5Ac7Ac to its 8-epimer (8ePse5Ac7Ac) in *A. baumannii* (29), possibly predicting 8ePse5Ac7Ac or a novel non-2-ulosonic acid isomer in the K264 CPS. The second locus, KL332, carried a novel gene named *qnlA*, which was found adjacent to a *rmlBDAC-rhnA* module. QnlA from KL332 shares 38.7% identity with QnlA from *Escherichia coli* O3 (GenPept accession number ABG81809.1) that is required for the synthesis of UDP-2,3-diacetamido-2,3,6-trideoxy-L-mannose (UDP-L-QuiNAc) (44). The third locus, KL363, carries *qdtF* and *qdtG* that replace *qdtC* and *qdtD* in the module of genes for synthesis of dTDP-3-acetamido-3,6-dideoxy-D-glucose (dTDP-D-Qui3NAc) (45), and the corresponding products share 63 or 70% amino acid identity, predicting either D-Qui3NAc or a potentially new variant of this sugar in K363.

### Frequency and distribution of genes at the K locus

To visualise patterns in genetic organisation, a gene presence/absence matrix was generated using homology groups defined at 85% translated protein identity (Supplementary Table 5). The seven conserved core genes, *wza, wzb, wzc, galU, ugd, gpi*, and *pgm,* that are essential for CPS biogenesis were present in all 409 K loci. Most were represented by three to five homology groups, though each formed a single cluster when the identity threshold was reduced to 60%. The exception was *pgm*, which was represented by a single homology group across all thresholds tested. These genes were always localised to the ends of each locus in the same orientation and modular order as shown in Fig. 1a, flanking Region 2 that carried varying combinations of type-specific structural genes. However, variation was also observed in Region 3 with many KL (n=373; 92.1%) including one or more genes between *gpi* and *pgm.* The presence of *gne1* at this location was common (Table 1), found either on its own (n=165 KL; 40.3%), adjacent to *pgt1* (n=146; 35.7%), or in combination with other genes (n=30; 7.3%). Hence, all novel loci exhibited the same overall genetic arrangement as described previously (17–19).

**Table 1.**
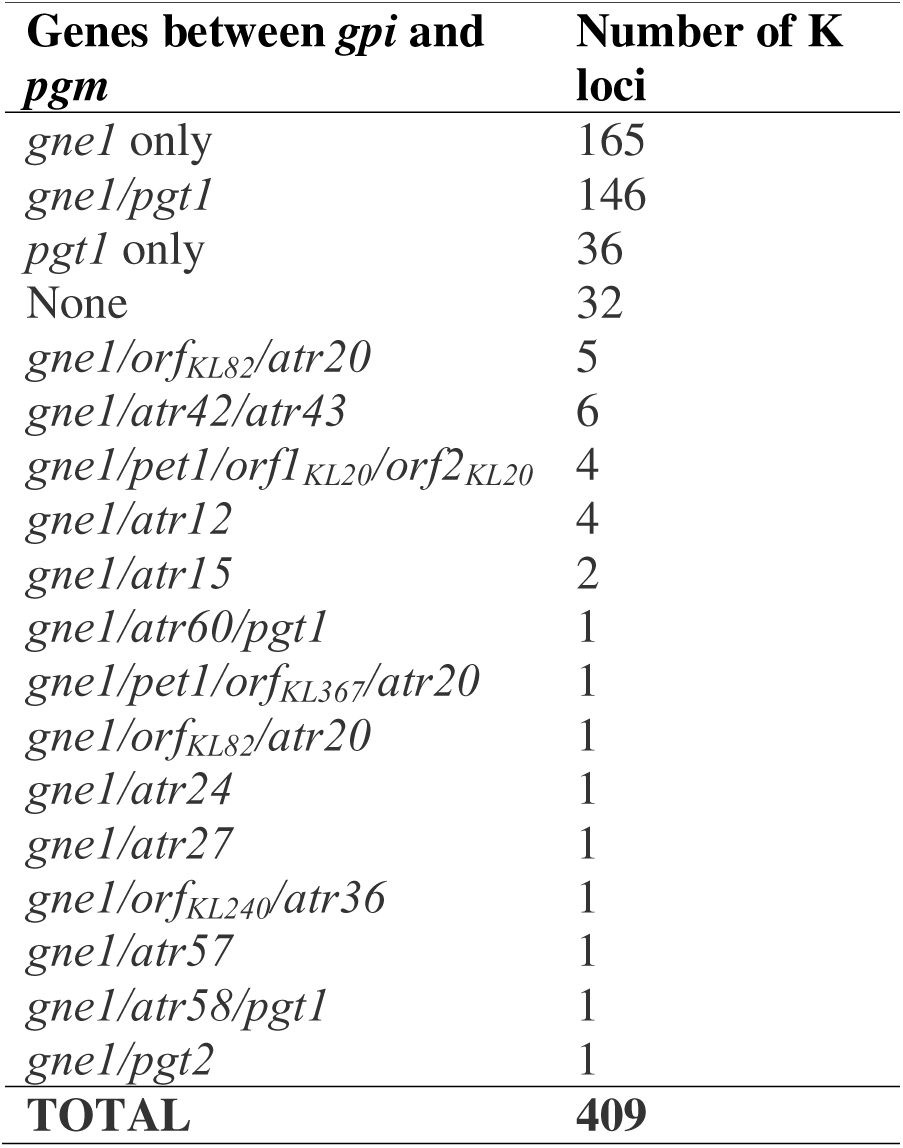
Number of KL and K types per combinations of genes found in Region 3.

In addition to conserved core genes, all loci harboured at least one gene for an initiating transferase (Itr), as well as for a Wzx translocase and Wzy polymerase that are also each essential for CPS biosynthesis in *A. baumannii* (19). The only exceptions were KL19, KL24 and KL39 that lack a *wzy* gene within the K locus but these KL co-occur with GI1 or GI2 that provide the Wzy polymerase necessary for CPS biogenesis (22, 23). Each locus also included a *gna* gene for a UDP-N-acetyl-D-glucosamine (UDP-D-GlcNAc) dehydrogenase, which was consistently located at the 5’ end of Region 2 (left hand side in Fig. 1a). Previously, three genes for homologues of Gna had been described at this location (17, 19) and no new variants were identified here. The *gna1* gene was found in 31 KL, 27 of which included *gne2* immediately downstream for production of UDP-D-GalNAc. The *gna3* gene in 46 KL always co-occurred with *dgaABC,* and the combination of these genes with *gna3* is responsible for UDP-2,3-diacetamido-2,3-dideoxy-α-D-glucuronic acid (UDP-D-GlcNAc3NAcA) synthesis as described previously (19). However, *gna2* was not associated with any specific genes. While its role in CPS biosynthesis has not been established (17, 19), it was the most common type-specific gene occurring in 332 KL (Fig. 3a).

**Figure 3.**
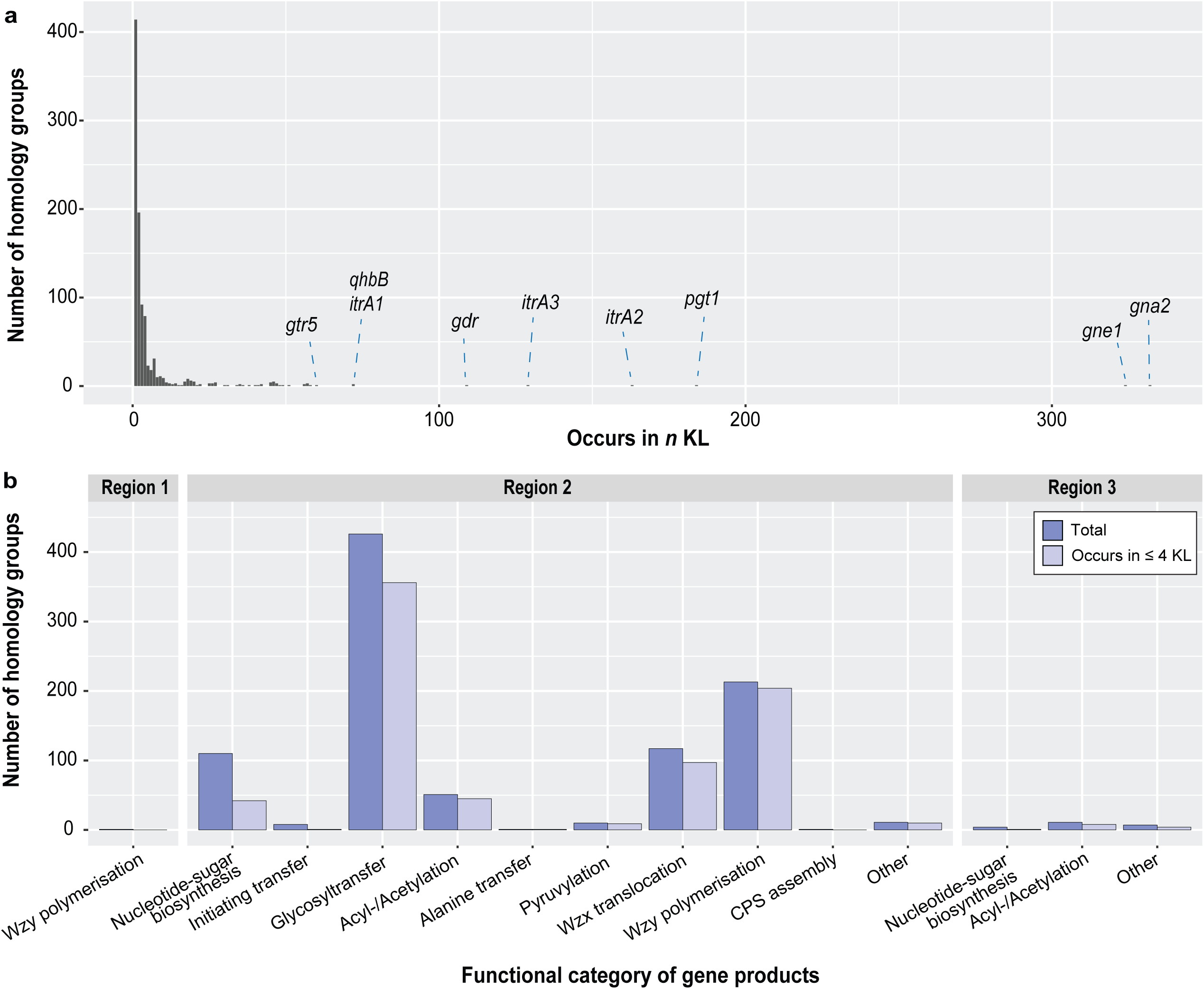
Distribution of variable genes across K loci. **(a)** Occurrence of individual genes across 409 K loci calculated from homology groups (85% translated identity threshold). Genes found in >60 KL are indicated. **(b)** Distribution of genes per functional category across each K locus region. Number of genes shared by ≤ 4 KL per category are shown in light blue. Functional categories assigned using the *A. baumannii* CPS gene nomenclature scheme (17–19).

Distribution of type-specific genes across Regions and functional categories (Fig. 3b) showed that the majority of clusters were from Region 2 and were those responsible for forming intra-unit glycosidic linkages (glycosyltransferases; n=429 clusters) or inter-unit glycosidic linkages (Wzy polymerases; n= 213), suggesting substantial diversity in CPS configuration and topology in *A. baumannii* as suggested previously (17). Wzx translocases also exhibited a high level of diversity with 117 clusters. Comparatively, decorative pyruvyl, O-acetyl or D-alanine groups were less common, with 123 loci encoding ≥1 Atr acetyl-/acyltransferase that were not associated with specific sugar biosynthesis pathways (48 possible *atr* genes in Region 2 and 11 in Region 3), 25 KL encoding one of ten Ptr pyruvyltransferases, and only 3 KL encoding Alt1 for the addition of D-alanine. There were 114 genes for complex sugar biosynthesis found in Region 2, with over half of them located in a defined module to the right of the *gna* gene (as shown in Fig. 1a). Visual representations of all K loci separated into groups based on gene(s) adjacent to *gna* are provided in Supplementary Figures 2-12.

Most type-specific structural genes (80.1%; n=781 of 975) were found in ≤ 4 loci, with a substantial portion of the total pool (42.5%; n=414 of 975) occurring in only a single KL (Fig. 3a). This is in line with the findings of a previous analysis on the genetic repertoire of 237 KL (17) and continues to suggest significant diversity in CPS types produced by *A. baumannii*. However, to better understand the extent of diversity in genes required for the determination of specific structural features (i.e. sugar constituents, intra-unit and inter-unit glycosidic linkages) predicted from all 409 loci, the frequency of these genes was reassessed in greater detail in the following sections.

### Genes for synthesis of nucleotide-linked sugar substrates

All K loci include genes for the synthesis of UDP-D-Glc and UDP-D-GlcNAc in Region 3, and these substrates can be converted to UDP-D-Gal or UDP-D-GalNAc when *gne1* is present (19). Across the 409 loci, most KL (n=359; 87.8%) also harboured genes(s) in Region 2 for synthesis of at least one other nucleotide-linked sugar. However, a total of 36 different gene modules were found (Fig. 4a), three of which are described here for the first time (see above). Genes for the synthesis of a non-2-ulosonic acid residue were present in 104 KL (25.4%; Fig. 4a), predicting either acetylated and/or acylated variants of pseudaminic acid (n=56 KL), legionaminic acid (n=26 KL), 8-epilegionaminic acid (n=2 KL), acinetaminic acid (n=6 KL), 8-epiacinetaminic acid (n=1 KL), neuraminic acid (n=2 KL) or two potentially novel isomers (n=11 KL). Genes for acetylated and/or acylated forms of bacillosamine (D-QuiNAc4NR) were in the next largest number of KL (n=72 KL), followed by genes for L-rhamnose (L-Rha; n=62 KL), N-acetyl-L-fucosamine (L-FucNAc; n=49 KL), N-acetyl-D-mannosaminuronic acid (D-ManNAcA; n=45) and D-GlcNAc3NAcA (n=46 KL). Genes predicting L-QuiNAc (*qnlA*), D-galactofuranose (D-Gal*f*; *glf*) or a possible variant of D-Qui3NAc (*qdtF-qdtG*) were rare, found in only one KL each.

**Figure 4.**
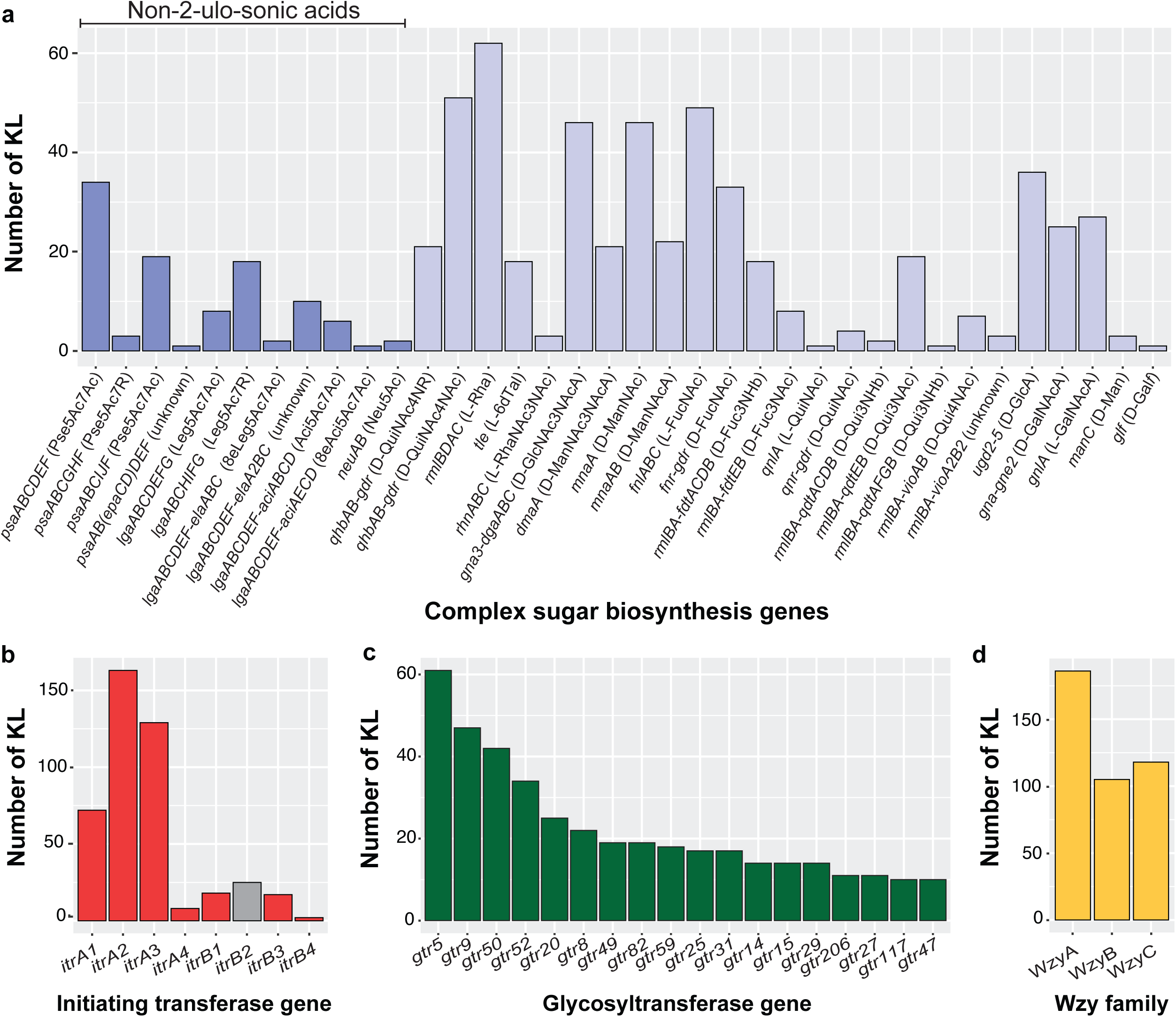
Frequency of genes responsible for determining CPS composition and topology. **(a)** Number of KL with genes in Region 2 responsible for complex sugar biosynthesis. Gene(s) predicting non-2-ulosonic acid isomers are indicated in dark blue. **(b)** Number of KL with genes encoding different Itr types. **(c)** Number of KL with glycosyltransferase genes that occur in more than 10 loci each. **(d)** Number of KL with genes encoding different Wzy family types as defined in (38).

### Genes for initiation of CPS biosynthesis

Construction of CPS units in *A. baumannii* begins with an initiating transferase (Itr) that forms a phosphodiester bond between a cytoplasmic membrane lipid carrier and one of six possible ‘first’ sugars. There are seven known Itr types in *A. baumannii* that fall into ItrA and ItrB superfamilies (19), though ItrB2 is reported to be redundant (19, 46). Each of the 409 KL encode at least one of these Itr types (Fig. 4b), and over half include either *itrA2* (n=163) or *itrA3* (n=129) specific for D-GalNAc (47, 48) and D-GlcNAc (49, 50), respectively. The *itrA1* gene is present in 72 KL and is always found with UDP-D-Qui*p*NAc4NR genes to provide its substrate (51). All other *itr* genes occur less frequently (Fig. 4b), including a gene for a fourth ItrB type that is identified in this study. The *itrB4* gene was found in KL385, and clustered with the *itr* gene from KL75 at the 85% aa threshold leading to the reassignment of the *itr* gene in KL75 as *itrB4*. ItrB4 is 81.6% identical to ItrB3, suggesting that it is a variant of ItrB3 and is likely to have specificity for the same or similar substrates. Finally, the *itrB2* gene is found only in a small number of KL that also include an *itrA2* (n=2) or *itrA3* (n=23) gene immediately adjacent to it.

### Genes for the formation of intra-unit glycosidic linkages

A total of 429 glycosyltransferase genes (*gtr*=424, *kpsS*=5) were found across the 409 loci, including 153 novel genes not reported previously (17). More than half of the encoded products (n=295) could be classified into CAZy families defining the mechanism (inverting or retaining) that determines the anomeric configuration of the glycosidic linkage formed. GT2 (n=108 Gtrs; inverting) and GT4 (n=135 Gtrs; retaining) families were common (Supplementary Table 3). However, many (n=134) could not be assigned to any established CAZy family, suggesting that these may represent new CAZy families as suggested for some Gtrs previously (e.g. 52, 53). The most common *gtr* genes were *gtr5* (n=61), *gtr9* (n=47), *gtr50* (n=42), *gtr52* (n=34) and *gtr20* (n=25) (Fig. 4c). These genes were mutually exclusive, collectively occurring in a total of 209 KL (51.1%). In solved CPS structures, all are known to be responsible for forming the first intra-unit linkage of the respective KUs, and all but *gtr20* are located immediately upstream of the *itr* gene. For *gtr20,* the *qnr3*-*itrB2* genes that both predict products reported to be redundant (19, 21, 46), are positioned between *gtr20* and the *itr* gene.

Previous studies have reported the prevalence of *gtr5, gtr9, gtr20* and *gtr50* genes across multiple loci, noting common pairings with at least one other *gtr* and a specific *itr* gene (e.g. 19, 21, 46, 54-58). Here, *gtr5, gtr9,* and *gtr50* genes were each found to pair with multiple *gtr* genes, and in some cases multiple *itr* genes suggesting relaxed specificity for the first sugar. Modules that include one of the five common *gtr* genes and are found in more than 8 loci are listed in Table 2, along with the linkages that they are known or predicted to produce. As studies have reported evidence that Gtr5 and Gtr50 may have relaxed substrate specificity (53, 58), further work will be needed to correlate amino acid differences with substrate and linkage specificities.

**Table 2.** Gene modules that include common *gtr* genes.

| Gene module | No.<br>KL | Structural segment | Reference |
| --- | --- | --- | --- |
| *gtr25-gtr5-itrA2* | 16 | α-D-Galp-(1→6)-β-D-Galp-(1→3)-D-GalpNAc | (45, 56, 72-74) |
| *gtr8-gtr9-itrA2* | 22 | α-D-Galp-(1→6)-β-D-Glcp-(1→3)-D-GalpNAc | (26, 27, 54, 56, 70-71) |
| *gtr77-gtr9-itrA2* | 8 | α-D-Galp-(1→6)-β-D-Glcp-(1→3)-D-GalpNAc | (55) |
| *gtr20-(qnr3-itrB3)-itrA3* | 23 | α-L-FucpNAc-(1→3)-D-GlcpNAc | (21, 46) |
| *gtr49-gtr50-itrA3* | 16 | α-D-GlcpNAc-(1→4)-α-D-GalpNAc-(1→3)-D-GlcpNAc | (23, 53) |
| *gtr117-gtr50-itrA3* | 8 | α-D-GalpNAc-(1→4)-α-D-GalpNAc-(1→3)-D-GlcpNAc | - |
| *gtr52-itrA1* | 36 | β-D-ManpNAcA-(1→3)-D-QuipNAc4NR | (69) |

### Genes determining the glycosidic linkage between K units

Wzy polymerases process fully formed oligosaccharide K units (KUs) into CPS polymers prior to export and determine the specific glycosidic linkage formed between adjacent Kus (38). A *wzy* polymerase gene could be identified in 406 K loci, the exceptions being KL19, KL24 and KL39 as described above. Three loci, KL67, KL134 and KL331, were unusual in that they each had two *wzy* genes, suggesting that there may be two different linkages between KUs in CPS of isolates with these loci. A small number of *wzy* genes (n=9) were shared by >4 loci (Fig. 3b), with genes coding for Wzy_KL2_ (n=10 KL), Wzy_KL60_ (n=10 KL), Wzy_KL9_ (n=7 KL), Wzy_KL5_ (n=6 KL), and Wzy_KL8_ (n=6 KL) the most common.

All KL-encoded Wzy included 8-14 TMS and could be classified into either WzyA, WzyB or WzyC protein families (Fig. 4d). These families have been shown to correlate with the anomeric configuration of the linkage formed (38), and as WzyA and WzyC predict β-configured linkages, the combined number of genes coding for WzyA and WzyC types across 293 of 406 loci indicate a preference for β-linkages between units in *A. baumannii* CPS as noted in a recent study (38).

### Several K loci predict the same CPS or KU

Previous studies have documented related K loci that either produce the same CPS (e.g. 25) or a CPS made up of the same KU (e.g. 59, 60). In reported cases of shared CPS, the structures are derived from loci that differ only in the presence of genes between *gpi* and *pgm* in Region 3, and a total of 23 groups of KLs that differ in this region have been reported previously (17). For loci that produce CPS made up of the same KUs, the loci differ in the type of *wzy* gene present. To uncover further locus groupings that predict the same CPS or KU, a dissimilarity matrix for unsupervised hierarchal clustering was computed for the 409 loci based on genes present in Region 2 only. Following clustering, the resulting dendrogram was plotted against a heatmap reporting gene presence (Fig. 5), which revealed several large clades that were generally associated with genes found at the 5’ end of Region 2 (left hand side in Fig. 1a). However, assessment of shorter branches of the dendrogram revealed several groups or pairs of KL that share the same or similar type-specific gene content.

**Figure 5.**
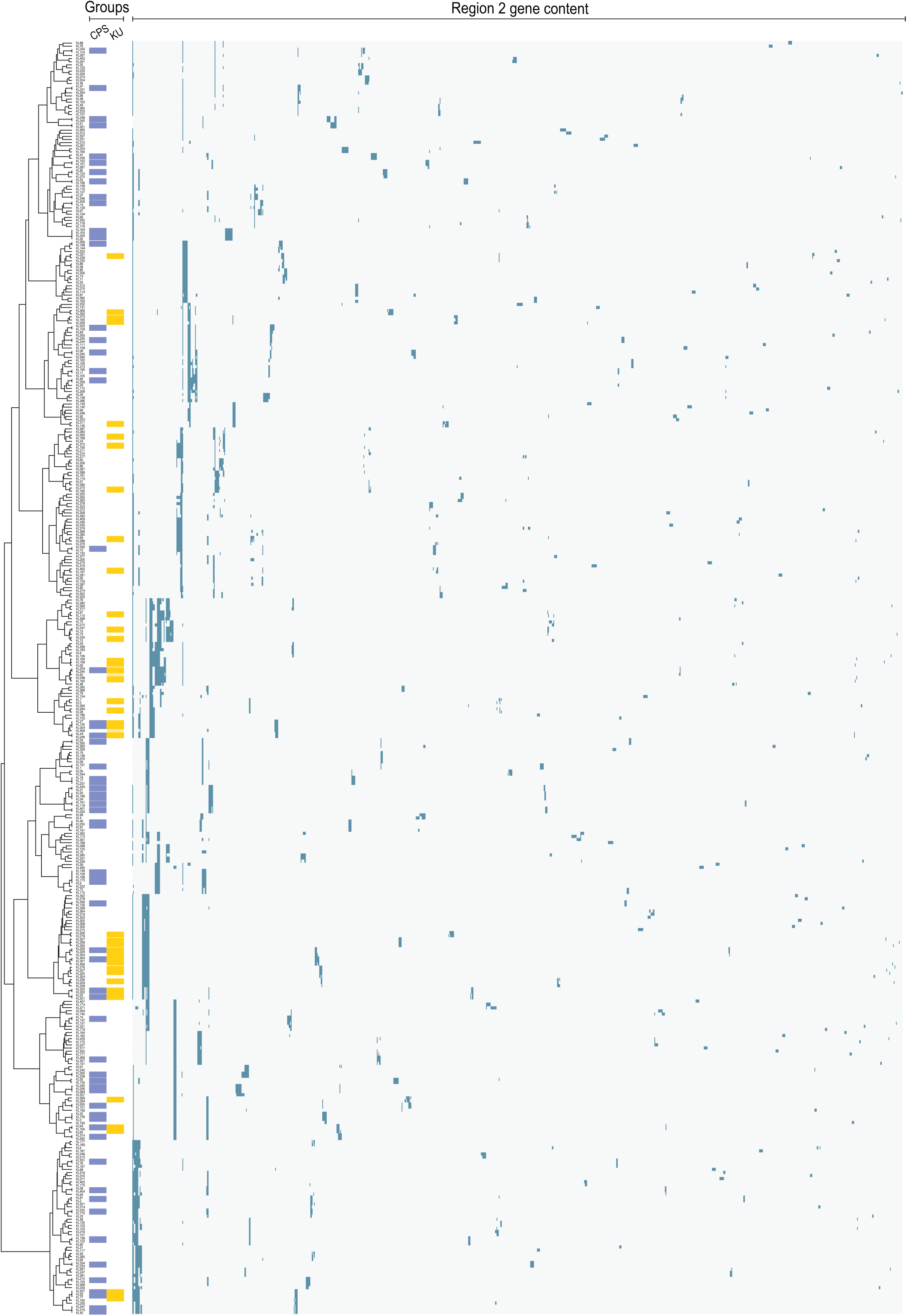
Hierarchal clustering of KL based on gene presence/absence. Dendrogram (left) constructed using the ward.D2 method determined as the optimal clustering algorithm using Region 2 homology groups calculated at 85% translated identity threshold. Heatmap of gene presence (green) and absence (grey is right. Blue column is grouped loci differing only in Region 3 that predict the same CPS. Yellow column Is grouped loci differing only in *wzy* that predict the same K unit (KU).

Inspection of these shorter branches found 119 KL that fell into 53 groups, which represented loci that differ in the region between *gpi* and *pgm* (blue groups in Fig. 5) extending upon the Region 3 groups reported previously (17). All 53 groups were further assessed for the types of genes found at this location (Supplementary Fig. 13). However, besides *gne1*, a role for these genes in CPS biosynthesis has not been established, indicating that members of these groups may produce the same CPS structure. Further inspection of the dendrogram led to the identification of 71 KL across 25 groups that represented loci that differed in the type of *w*zy polymerase gene present (yellow groups in Fig. 5). This finding suggests the occurrence of several CPS types that are made up from the same KU with a different linkage between units. In addition to these groupings, there were several other KL found to differ by 1-2 genes associated with sugar biosynthesis, glycosyltransfer or initiating transfer, indicating that many CPS structures produced by *A. baumannii* may be closely related.

### Frequency of K loci amongst sequenced *A. baumannii* isolates

Curated reference sequences for novel KL were incorporated into the *Kaptive* database to produce an updated version that comprises 409 KL and 10 extra-locus genes. To validate the utility of this database, the pool of 46,185 genome assemblies was rescreened using *Kaptive* with the same parameters (Supplementary Table 6) and the frequency of KL amongst publicly available *A. baumannii* sequences was assessed. Of the genomes that had a detectable K locus, 43,822 (94.9%) had a ‘Typeable’ categorical confidence level (Table 3), including 7,995 with fragmented matches and 42 with evidence of low sequence/assembly quality (see above). For matches considered ‘Untypeable’ (n=2,356; 5.1%), manual sequence inspection revealed that 447 were incorrectly assigned, many of which were due to allelic exchange of conserved core genes with other loci. Other ‘Untypeable’ assignments included 1,846 genomes with fragmented KL that were not examined in the study. Hence, a total of 44,227 (95.8%) genomes were assigned matches to a KL reference sequence in the updated database (Fig. 6a), indicating a high genome coverage.

**Figure 6.**
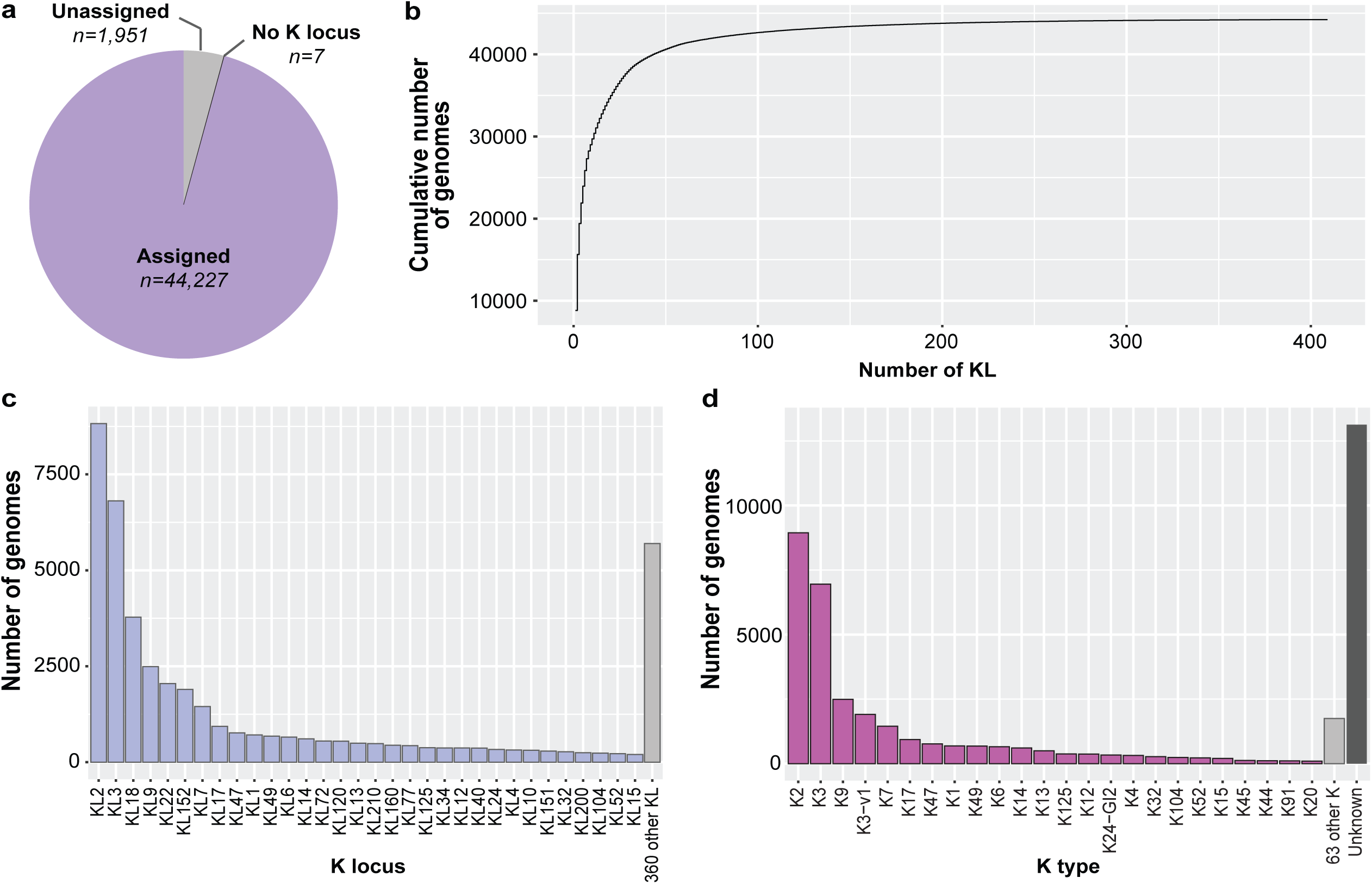
Distribution of K loci and K types predicted from *A. baumannii* genomes available in NCBI. **(a)** Proportion of genomes with high-confidence assignments made by Kaptive or via manual inspection. **(b)** KL coverage in available genomes with assigned K loci. **(c)** Proportion of assigned genomes carrying KL found in >200 genomes each. **(d)** Proportion of assigned genomes predicting distinct K types with available structural data.

**Table 3.**
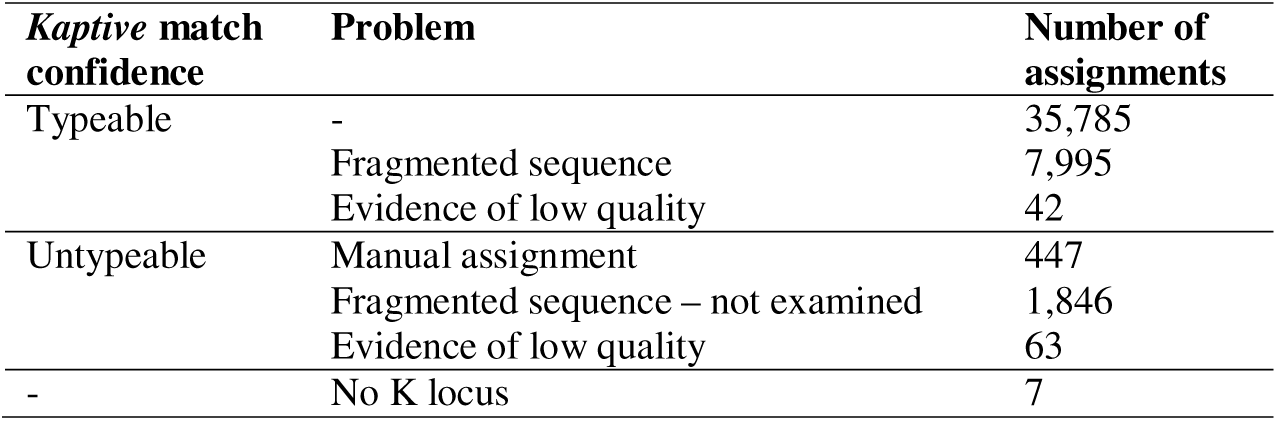
Summary of results for validation of the updated database against 46,185 *A. baumannii* genomes.

The total frequency of KL types amongst the 44,227 genomes (Fig. 6b) revealed that 17 KL were not found in the genome dataset and 75 KL were in only a single genome. Most novel loci reported in this study were found in ≤10 genomes each. KL258 (n=53), KL285 (n=40), KL309 (n=141) and KL370 (n=58) were the most common, with KL258 and KL285 notably associated with multiple countries and isolation sources. Only 32 KL occurred in >200 genomes, and these collectively represented 87.1% (n=38,530) of genomes with assigned KLs (Fig. 6c). However, over half carried either KL2 (n= 8,823), KL3 (n=6,809), KL18 (n= 3,778), KL9 (n=2,491), KL22 (n=2,048) or KL152 (n=1,898).

Granted *Kaptive* can infer K type based on the combination of KL and extra-locus genes where structural data is available, the frequency of predicted K types was also assessed. There were 87 K types predicted, 24 of which were represented by >100 genomes each (Fig. 6d). Of the 8,857 genomes with either KL3 or KL22, 6,951 predict the K3 type whereas 1,906 had a detectable *gtr6* truncation predicting the K3-v1 type. While many genomes (n=13,129) carried KL with no currently associated structural data, a large proportion had KL18 (n= 3,778) that is a Region 3 pair with KL17 (Supplementary Fig. 13). While there is no current structural information available for a KL18 isolate, it has been previously predicted to produce the K17 CPS (61).

### Frequency of extra-locus genes in sequenced *A. baumannii* isolates

Granted their importance to CPS typing, the frequency of the 10 documented extra-locus genes (from GIs and prophage) in the genome dataset was also examined. A copy of GI1 was detected in 152 assemblies (Table 4), that harboured either KL1 (n=22), KL19 (n=82) or KL39 (n=48). It was also found on a short contig (7114 bp) in an assembly that included KL12, though the boundaries of the island could not be identified. In addition, *wzyGI1* was detected in three assemblies with KL146 (isolates named 1404.312, AB053 and AB048; NCBI accession numbers listed in Supplementary Table S1), one of which (isolate 1404.312) also carried *atr25.* A comparison of KL146 with KL1, KL19 and KL39 is shown in Fig. 7a and demonstrates that KL146 is closely related to KL19. These two loci share the same type-specific genes responsible for production of a shared KU, with KL19 likely arising from KL146 via deletion of the *wzy_KL146_* and *atr31* genes. An insertion adjacent to *cpn60* was not found in any of the three KL146 genomes, and inspection of the genetic context of *wzyGI1* and/or *atr25* revealed that these genes were in one of two distinct prophages in these genomes (Fig. 7b). The encoded proteins were not identical to those from GI1, suggesting a complex evolutionary history.

**Figure 7.**
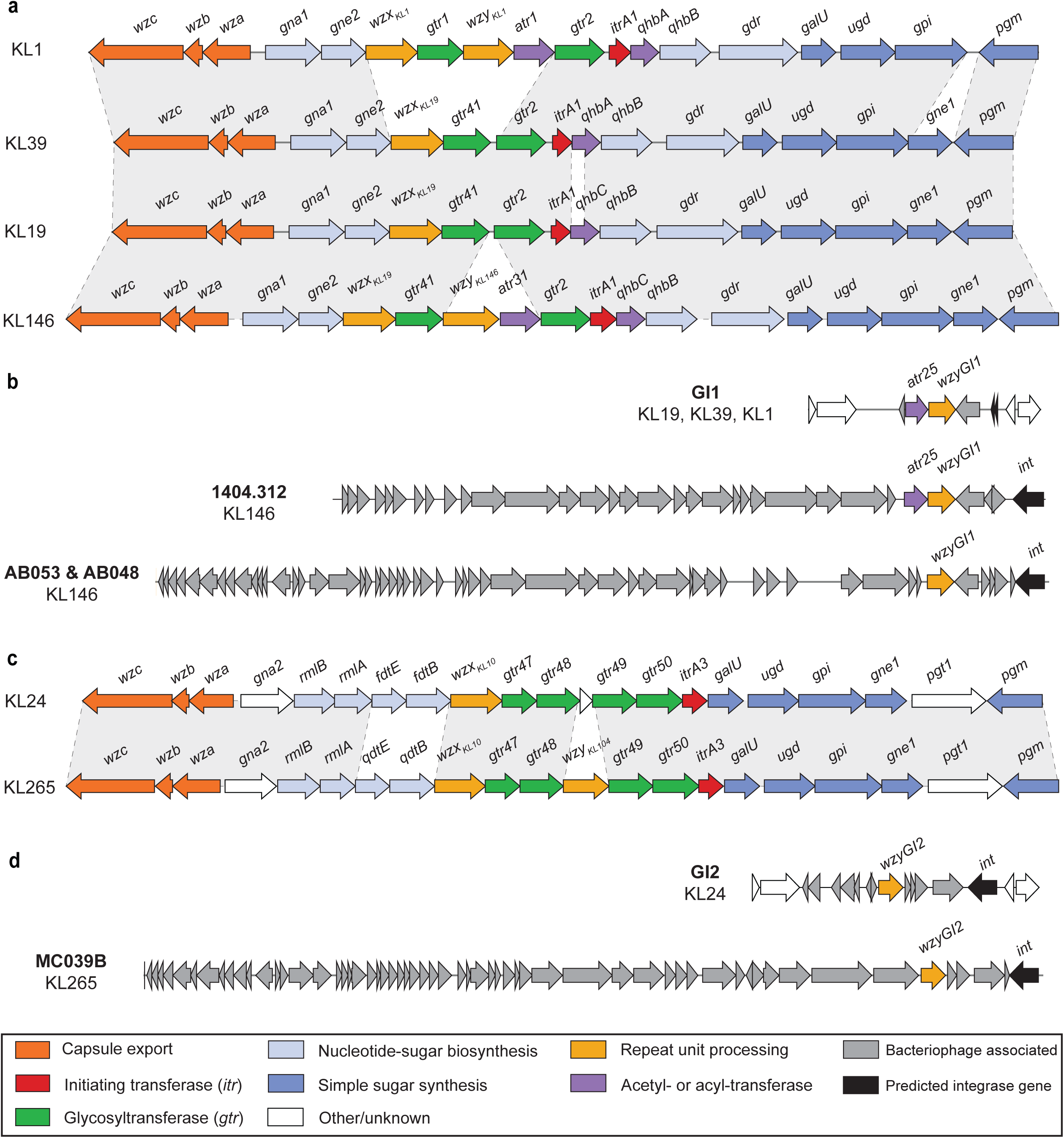
K loci and prophage with CPS biosynthesis genes found in isolates with *wzyGI1* or *wzyGI2* but without a known genomic island. (a) Comparison of KL19 and KL146 associated with *wzyGI1.* (b) Comparison of GI1 with prophage that include *wzyGI1.* (c) Comparison of KL24 and KL265 associated with *wzyGI2.* (d) Comparison of GI2 with prophage that include *wzyGI2.* Colour scheme for genes in below, and grey shading between KL indicates segments with shared genes. Figure was constructed in *pygenomeviz v 1.6.1* (https://github.com/moshi4/pyGenomeViz) and annotated in Adobe Illustrator.

**Table 4.** **Summary of known extra-locus genes detected in sequenced *A. baumannii***

| Extra-locus gene(s) | Genetic context | K loci (n) | Total no. of genomes |
| --- | --- | --- | --- |
| *atr25* + *wzyGI1* | GI1 | KL1 (22), KL12 (1), KL19 (80), KL39 (48) | 151 |
| *wzyGI1* | GI1 | KL19 (2) | 2<sup>1</sup> |
| *wzyGI2* | GI2 | KL24 (322) | 322 |
| *atr25* + *wzyGI1* | Prophage | KL146 (1) | 1 |
| *wzyGI1* | Prophage | KL146 (2) | 2 |
| *wzyGI2* | Prophage | KL265 (1) | 1 |
| *wzy1Ph* | Prophage | KL127 (24) | 24 |
| *wzy2Ph* | Prophage | KL41 (2), KL58 (5), KL79 (11), KL94 (4), KL170 (1), KL225 (2), KL259 (1), KL316 (1) | 27 |
| *wzy1Ph* + *wzy2Ph* | Prophage | KL58 (1) | 1 |
| *atr29* | Prophage | KL46 (4), KL120 (1) | 5 |
| *atr30* | Prophage | KL5 (5) | 5 |
| *atr44* | Prophage | KL58 (3), KL60 (2), KL155 (1), KL177 (2) | 8 |
| *epaA-epaB* | Prophage | KL30 (9), KL46 (1), KL58 (10), KL60 (2), KL80 (1), KL117 (2), KL120 (10), KL135 (3), KL334 (4), KL361 (3), KL381 (1) | 46 |
| *epaB* | Prophage | KL6 (4), KL23 (6), KL49 (3), KL247 (1) | 14<sup>2</sup> |
| *epaA-epaB* + *wzy2Ph* | Prophage | KL58 (1) | 1 |
| *epaA-epaB* + *atr44* | Prophage | KL117 (1) | 1 |

<sup>1</sup> *atr25* is truncated in these genomes
<sup>2</sup> *epaA* is truncated in these genomes

A complete copy of GI2 was detected in all 322 genomes that harboured KL24 (Table 4). However, *wzyGI2* was also found in a single genome with KL265 (isolate MC039B; NCBI accession number GCA_045158015.1). KL24 and KL265 (Fig. 7c) were also found to be closely related, sharing similar type-specific genes. However, KL265 included *wzy_KL104_* that was absent in KL24 and had a module of genes for D-Qui3NAc in place of the genes for D-Fuc3NAc. Similar to the situation for the three KL146 genomes described above, there was no detectable insertion at the *cpn60* locus and *wzyGI2* was found in a prophage (Fig. 7d). As KL146 and KL265 both carry a *wzy* polymerase gene that appears to predict a functional product, the *wzyGI1* and *wzyGI2* genes in prophage may be involved in modifying the linkage between KUs as in the case for other isolates reported with prophage-encoded Wzy (30, 31).

In addition to genes in GI1 and GI2, there are three types of extra-locus genes relevant to CPS typing that are found in prophage. These include those that either: (i) change sugars in select CPS structures; (ii) alter the specific linkage between oligosaccharide KUs in the CPS; or (iii) decorate sugars in KUs with acetyl groups. Only 132 (0.3%) assemblies were found to include 1 or more of the known extra-locus genes from prophage (Table 4; presence defined by >85% translated protein sequence identity and >80% coverage). These were found in combination with one of 26 different KLs. While some of these combinations have been reported previously and have associated structural data (22, 23, 27–31, 62), several new combinations are reported here for the first time. The *wzy1Ph* and *wzy2Ph* genes were found in 53 genomes associated with a variety of KLs (Table 4) and were only found together in a single genome from isolate BAL114, consistent with the findings of a recent analysis on these genes (31). Genes for acetyltransferases, previously been shown to decorate K5 (*atr30*) and K46 (*atr29*) (28), and K58 (*atr44*) (27), were found in 20 genomes. Finally, genes for sugar biosynthesis enzymes, EpaA and EpaB, that convert Pse5Ac7Ac to 8ePse5Ac7Ac (29) were in 62 genomes. These carried 15 different KL, four of which are reported here for the first time and include *psa* genes for synthesis of the pseudaminic acid substrate.

## DISCUSSION

As a major protective cell-surface structure and essential virulence determinant, the CPS is subject to diversifying selection that gives rise to an extraordinary number of possible CPS types produced by a species (63). Prior to this study, a total of 241 distinct K loci had been officially documented for *A. baumannii* in the CPS typing database compatible with *Kaptive* (17). Here, the addition of 168 novel loci brings the total number of KL described for *A. baumannii* to 409, which far exceeds the number of capsule biosynthesis loci reported for other pathogens, including *Klebsiella pneumoniae* (64, 65)*, Escherichia coli* (66, 67), and *Streptococcus pneumoniae* (68). Though there are known cases of KL that produce the same CPS structure but differ in Region 3 necessitating the assignment of a distinct KL identifier (17), the consolidation of KL sequences that share type-specific genes still predicts more than 330 unique CPS structures from Region 2 genes alone. However, extra-locus genes, IS and frameshift mutations can contribute to further diversity that was not explored in detail in this study.

While all novel K loci reported in this study conform to the same locus configuration described previously (19), there were 309 protein clusters representing novel genes. The function of members of most clusters could be predicted using a combination of sequence-based approaches, and these predictions were verified via the screening of modelled tertiary structures against available PDB entries. This work highlights that a variety of tools is currently necessary annotate and assign functions to enzymes associated with CPS biosynthesis in *A. baumannii*. However, the curated set of fully annotated sequences available in the updated database, in addition to modelled tertiary structures for members of all 1010 protein clusters, will serve as a valuable and scalable resource for CPS gene identification and sequence annotation into the future.

Analysis of gene distribution by functional category revealed significant diversity in genes responsible for the determination of CPS type, including an extensive repertoire of genes predicting 36 different sugars and >420 intra-unit (Gtr/KpsS) and >200 inter-unit (Wzy) glycosidic linkages. Despite this diversity, hierarchal clustering of KL based on Region 2 gene content revealed many groups of closely related loci predicting shared or similar CPS structures. Assessment of type-specific genes also revealed shared structural features, including the presence of genes for synthesis of sugars belonging to the 5,7-diamino-3,5,7,9-tetradeoxy-non-2-ulosonic acids subclass of complex non-2-ulosonic acids in 104 KL. The distribution of these genes, and those predicting other common sugars, will be useful for informing the generation of further monoclonal antibodies specific for non-2-ulosonic acid CPS constituents (7, 8). However, the most common gene outside of the conserved core CPS genes was *gna2*. As this gene encodes a predicted UDP-D-GlcNAc dehydrogenase that has no documented role in CPS biosynthesis (19), further work will be needed to understand its function and the reasons why it is present in the K locus.

Curated annotations for novel loci were integrated into the *Kaptive* database, which was then validated by rescreening the pool of 46,185 genomes. Only a small fraction of these were misassigned by *Kaptive*, and in most cases, misassignments were due to closer matches of conserved core genes with other reference sequences. Users of the updated database are therefore encouraged to inspect raw outputs to confirm typing results. Here, 87.1% of studied genomes accounted for only 32 KL, with overrepresented sequences including KL2, KL3, KL9, KL18, and KL22. However, known sampling bias in public datasets associated with large regional or clonal studies, as observed with the larger number of genomes from the USA and China in the dataset utilised in this study (Supplementary Fig. 1a), restricts further exploration of CPS prevalence, temporal shifts and the association with antibiotic resistance, and dereplication of the dataset was beyond the scope of this study. Nonetheless, there has been an evident expansion of *A. baumannii* genome sequences released into NCBI from diverse geographical locations and isolation sources (Supplementary Fig. 1), including non-human sources that are suggested to have an open pan genome (3). Therefore, the expansion of the CPS typing infrastructure for *A. baumannii* will greatly assist with ongoing epidemiological investigations and strain characterisation.

Several studies have now reported the presence of extra-locus genes in *A. baumannii* genomes that influence the composition and topology of the CPS (27–31). GI1 and GI2 remain the only characterised genomic islands that include genes relevant to CPS typing. However, homologues of the extra-locus genes that they carry were found here in prophage, indicating that GI1 and GI2 may have a complex evolutionary history and may also be associated with bacteriophage elements. Detection of CPS genes outside the K locus has historically been challenging due to a lack of tools and available genotype-structure maps for individual isolates (38). Hence, it is possible that the extent of extra-locus genes that influence the CPS type may be underappreciated in *A. baumannii*, and the true level of CPS diversity likely higher than anticipated based on K locus diversity. As further extra-locus genes are discovered, the addition of these sequences and ongoing curation of the database will enable a deeper analysis of the frequency of extra-locus genes and the extent of their role(s) in generating CPS diversity.

## ACKNOWLEDGEMENTS

Ruth Hall (University of Sydney, Australia) is thanked for valuable discussions regarding updates to the *A. baumannii* CPS nomenclature and typing scheme and for critical reading of this manuscript. Kelly Wyres and Thomas Stanton (Monash University, Australia) are also thanked for database migration and continued discussions relating to updates relevant to this study. In addition, Francios Lebreton and Patrick McGann (Walter Reed Army Institute of Research (WRAIR), USA) are acknowledged for providing access to sequences with novel KL for release into public databases.

## AUTHOR CONTRIBUTIONS

Conceptualisation: JJK

Data curation: JJK

Formal analysis: JJK

Funding acquisition: JJK

Methodology: JJK

Project administration: JJK

Validation: JJK

Visualisation: JJK

Writing - original draft: JJK

Writing - revising and editing: JJK

## REFERENCES

1. Murray CJL, Ikuta KS, Sharara F, Swetschinski L, Aguilar GR, Gray A, et al. Global burden of bacterial antimicrobial resistance in 2019: a systematic analysis. Lancet. 2022;399(10326):629–55.

2. Sati H, Carrara E, Savoldi A, Hansen P, Garlasco J, Campagnaro E, et al. The WHO Bacterial Priority Pathogens List 2024: a prioritisation study to guide research, development, and public health strategies against antimicrobial resistance. Lancet: Infect Dis. 2024;S1473–3099(25):00118-5.

3. Aguilar-Vera A, López-Sánchez R, Hernández-Alvarez AJ, Vargas-Peralta D, Peralta H, Castillo-Ramírez S. Global multi-host genomic epidemiology of *Acinetobacter baumannii* reveals transmission at one health interfaces. Nat Commun. 2026;17:5832.

4. Li S, Jiang G, Wang S, Wang M, Wu Y, Zhang J, et al. Emergence and global spread of a dominant multidrug-resistant clade within *Acinetobacter baumannii*. Nat Commun. 2025;16(1):2787.

5. Castillo-Ramírez S, Aguilar-Vera A, Kumar A, Evans B. *Acinetobacter baumannii:* much more than a human pathogen. Antimicrob Agents Chemother. 2025;69(9):e00801–25.

6. Yang N, Jin X, Zhu C, Gao F, Weng Z, Du X, et al. Subunit vaccines for *Acinetobacter baumannii*. Front Immunol. 2023;13:1088130.

7. Tang AH, Soler NM, Karlic KI, Corcilius L, Clarke-Shepperson CE, Lehmann C, et al. Uncovering bacterial pseudaminylation with pan-specific antibody tools. Nat Chem Biol. 2026:1–12.

8. Slarve M, Jaramillo H, Luna B, Spellberg B. A monoclonal antibody raised against *Acinetobacter baumannii* capsular carbohydrate exhibits cross-species in vitro binding against Pseudomonas aeruginosa. PLoS One. 2026;21(1):p.e0340857.

9. Nielsen TB, Yan J, Slarve M, Lu P, Li R, Ruiz J, et al. Monoclonal antibody therapy against *Acinetobacter baumannii*. Infect Immun. 2021;89(10):e0016221.

10. Soontarach R, Srimanote P, Enright MC, Blundell-Hunter G, Dorman MJ, Thomson NR, et al. Isolation and characterisation of bacteriophage selective for key *Acinetobacter baumannii* capsule chemotypes. Pharmaceuticals. 2022;15(4):443.

11. Oliveira H, Domingues R, Evans B, Sutton JM, Adriaenssens EM, Turner DT. Genomic diversity of bacteriophages infecting the genus *Acinetobacter*. Viruses. 2022;14(2):181.

12. Rao S, Betancourt-Garcia M, Kare-Opaneye YO, Swierczewski BE, Bennett JW, Horne BA, et al. Critically ill patient with multidrug-resistant *Acinetobacter baumannii* respiratory infection successfully treated with intravenous and nebulized bacteriophage therapy. Antimicrob Agents Chemother. 2021;66(1):e00824–21.

13. Gordillo Altamirano F, Forsyth JH, Patwa R, Kostoulias X, Trim M, Subedi D, et al. Bacteriophage-resistant *Acinetobacter baumannii* are resensitized to antimicrobials. Nat Microbiol. 2021;6(2):157–61.

14. Schooley R, Biswas B, Gill J, Hernandez-Morales A, Lancaster J, Lessor L, et al. Development and use of personalized bacteriophage-based therapeutic cocktails to treat a patient with a disseminated resistant *Acinetobacter baumannii* infection. Antimicrob Agents Chemother. 2017;61:e00954–17.

15. Knirel YA, Shneider MM, Popova AV, Kasimova AA, Senchenkova SN, Shashkov AS, et al. Mechanisms of *Acinetobacter baumannii* capsular polysaccharide cleavage by phage depolymerases. Biochem (Mosc). 2020;85(5):567–74.

16. Popova AV, Shneider MM, Arbatsky NP, Kasimova AA, Senchenkova SN, Shashkov AS, et al. Specific interaction of novel Friunavirus phages encoding tailspike depolymerases with corresponding *Acinetobacter baumannii* capsular types. J Virol. 2021;95(5):e01714–20.

17. Cahill SM, Hall RM, Kenyon JJ. An update to the database for *Acinetobacter baumannii* capsular polysaccharide locus typing extends the extensive and diverse repertoire of genes found at and outside the K locus. Microb Genom. 2022;8:e000878.

18. Wyres KL, Cahill SM, Holt KE, Hall RM, Kenyon JJ. Identification of *Acinetobacter baumannii* loci for capsular polysaccharide (KL) and lipooligosaccharide outer core (OCL) synthesis in genome assemblies using curated reference databases compatible with Kaptive. Microb Genom. 2020;6(3):e000339.

19. Kenyon JJ, Hall RM. Variation in the complex carbohydrate biosynthesis loci of *Acinetobacter baumannii* genomes. PLoS One. 2013;8(4):e62160.

20. Kasimova AA, Sharar NS, Ambrose SJ, Knirel YA, Shneider MM, Timoshina OY, et al. The *Acinetobacter baumannii* K70 and K9 capsular polysaccharides consist of related K-units linked by the same Wzy polymerase and cleaved by the same phage depolymerases. Microbiol Spectr. 2023;11(6):e03025–23.

21. Arbatsky NP, Kenyon JJ, Kasimova AA, Shashkov AS, Shneider MM, Popova AV, et al. K units of the K8 and K54 capsular polysaccharides produced by *Acinetobacter baumannii* BAL 097 and RCH52 have the same structure but contain different di-N-acyl derivatives of legionaminic acid and are linked differently. Carbohydr Res. 2019;483:107745.

22. Kenyon JJ, Shneider MM, Senchenkova SN, Shashkov AS, Siniagina MN, Malanin SY, et al. K19 capsular polysaccharide of *Acinetobacter baumannii* is produced via a Wzy polymerase encoded in a small genomic island rather than the KL19 capsule gene cluster. Microbiology. 2016;162(8):1479–89.

23. Kenyon JJ, Kasimova AA, Shneider MM, Shashkov AS, Arbatsky NP, Popova AV, et al. The KL24 gene cluster and a genomic island encoding a Wzy polymerase contribute genes needed for synthesis of the K24 capsular polysaccharide by the multiply antibiotic resistant *Acinetobacter baumannii* isolate RCH51. Microbiology. 2017;163:355–63.

24. Kenyon JJ, Marzaioli A, Hall RM, De Castro C. Structure of the K2 capsule associated with the KL2 gene cluster of *Acinetobacter baumannii*. Glycobiology. 2014;24(6):554–63.

25. Talyansky Y, Nielsen TB, Yan J, Carlino-Macdonald U, Di Venanzio G, Chakravorty S, et al. Capsule carbohydrate structure determines virulence in Acinetobacter baumannii PLoS Pathog. 2021;17:e1009291.

26. Timoshina OY, Kasimova AA, Shneider MM, Arbatsky NP, Shashkov AS, Shelenkov AA, et al. Loss of a branch sugar in the *Acinetobacter baumannii* K3-type capsular polysaccharide due to frameshifts in the *gtr6* glycosyltransferase gene leads to susceptibility to phage APK37.1. Microbiol Spectr. 2023;11(1):e0363122.

27. Iovine A, Filatov AV, Kasimova AA, Sharar NS, Ambrose SJ, Dmitrenok AS, et al. Structure of the K58 capsular polysaccharide produced by *Acinetobacter baumannii* isolate MRSN 31468 includes Pse5Ac7Ac that is 4-O-acetylated by a phage-encoded acetyltransferase. Carbohydr Res. 2025;547:109324.

28. Kenyon JJ, Arbatsky NP, Shneider MM, Popova AV, Dmitrenok AS, Kasimova AA, et al. The K46 and K5 capsular polysaccharides produced by *Acinetobacter baumannii* NIPH 329 and SDF have related structures and the side-chain non-ulosonic acids are 4-O-acetylated by phage-encoded O-acetyltransferases. PLoS One. 2019;14(6):e0218461.

29. Sharar NS, Iovine A, De Castro C, Hall RM, Kenyon JJ. Phage-encoded enzymes found in *Acinetobacter baumannii* convert pseudaminic acid to 8-epipseudaminic acid. Commun Biol. 2025;8:700.

30. Arbatsky NP, Kasimova AA, Shashkov AS, Shneider MM, Popova AV, Shagin DA, et al. Involvement of a phage-encoded Wzy protein in the polymerization of K127 units to form the capsular polysaccharide of *Acinetobacter baumannii* isolate 36-1454. Microbiol Spectr. 2022;27:e0150321.

31. Sharar NS, Harmer CJ, Hall RM, Kenyon JJ. Multiple prophage acquistion events over the course of an outbreak drive lysogenic conversion of capsular polysaccharides produced by carbapenem-resistant *Acinetobacter baumannii* isolates. Microbiol Spectr. 2026:e04084–25.

32. O’Leary NA, Cox E, Holmes JB, Anderson WR, Falk R, Hem V, et al. Exploring and retrieving sequence and metadata for species across the tree of life with NCBI Datasets. Sci Data. 2024;11(1):732.

33. Stanton TD, Hetland MAK, Löhr IH, Holt KE, Wyres KL. Fast and accurate *in silico* antigen typing with *Kaptive* 3. Microb Genom. 2025;11:001428.

34. Gautreau G, Bazin A, Gachet M, Planel R, Burlot L, Dubois M, et al. PPanGGOLiN: Depicting microbial diversity via a partitioned pangenome graph. PLOS Comput Biol. 2020;16(3):e1007732.

35. Potter S, Luciani A, Eddy S, Park Y, Lopez R, Finn R. HMMER web server: 2018 update. Nucleic Acids Res. 2018;46(Web Server Issue):W200–W4.

36. Pandurangan AP, Stahlhacke J, Oates ME, Smithers B, Gough J. The SUPERFAMILY 2.0 database: a significant proteome update and a new webserver. Nucleic Acids Res. 2019;47(D1):D490–D4.

37. Zheng J, Ge Q, Yan Y, Zhang X, Huang L, Yin Y. dbCAN3: automated carbohydrate-active enzyme and substrate annotation. Nucleic Acids Res. 2023;51(W1):W115–W21.

38. Stanton TD, Ulacco L, Hall RM, Wyres KL, Kenyon JJ. Redefining bacterial Wzy polymerase families via three-dimensional structure-based clustering. 2026. DOI: 10.21203/rs.3.rs-9286338/v1

39. Abramson J, Adler J, Dunger J, Evans R, Green T, Pritzel A, et al. Accurate structure prediction of biomolecular interactions with AlphaFold 3. Nature. 2024;630(8016):493–500.

40. Van Kempen M, Kim SS, Tumescheit C, Mirdita M, Lee J, Gilchrist CL, et al. Fast and accurate protein structure search with Foldseek. Nat Biotechnol. 2024;42(2):243–6.

41. Kassambara A. Practical guide to cluster analysis in R: Unsupervised machine learning: Sthda; 2017.

42. Wickham H. Data analysis. In ggplot2 Springer, Cham.; 2016.

43. Wishart DS, Han S, Saha S, Oler E, Peters H, Grant JR, et al. PHASTEST: faster than PHASTER, better than PHAST. Nucleic Acids Res. 2023;51(W1):W443–W50.

44. Liu B, Furevi A, Perepelov AV, Guo X, Cao H, Wang Q, et al. Structure and genetics of *Escherichia coli* O antigens. FEMS Microbiol Rev. 2019;44(6):655–83.

45. Arbatsky NP, Shashkov AS, Sharar NS, Baird FJ, Shneider MM, Shpirt AM, et al. The K95 capsular polysaccharide produced by *Acinetobacter baumannii* isolate MAR18-2212 includes a rarely encountered 3-acetamido-3, 6-dideoxy-D-glucose (D-Qui3NAc) sugar. Carbohydr Res. 2025;553:109499.

46. Kasimova AA, Shneider MM, Shashkov AS, Kenyon JJ, Hall RM, Knirel YA. Structure of the K49 capsular polysaccharide produced by the *Acinetobacter baumannii* ST10 carriage isolate, NL6. Carbohydr Res. 2025;557:109646.

47. Kenyon JJ, Marzaioli, A.M., Hall, R.M. and De Castro, C. Structure of the K6 capsular polysaccharide from *Acinetobacter baumannii* isolate RBH4. Carbohydr Res. 2015;409:30–5.

48. Lees-Miller RG, Iwashkiw JA, Scott NE, Seper A, Vinogradov E, Schild S, et al. A common pathway for *O*-linked protein-glycosylation and synthesis of capsule in *Acinetobacter baumannii*. Mol Microbiol. 2013;89(5):816–30.

49. Kenyon JJ, Shashkov AS, Senchenkova SN, Shneider MM, Liu B, Popova AV, et al. *Acinetobacter baumannii* K11 and K83 capsular polysaccharides havethe same 6-deoxy-l-talose-containing pentasaccharide K units butdifferent linkages between the K units. Int J Biol Macromol. 2017;103:648–55.

50. Harding C, Haurat F, Vinogradov E, Feldman M. Distinct amino acid residues confer one of three UDP-sugar substrate specificities in *Acinetobacter baumannii* PglC phosphoglycosyltransferases. Glycobiology. 2018;28(7):522–33.

51. Morrison MJ, Imperiali B. Biosynthesis of UDP-*N,N*’-diacetylbacillosamine in *Acinetobacter baumannii*: Biochemical characterization and correlation to existing pathways. Arch Biochem Biophys. 2013;536:72–80.

52. Kenyon JJ, Kasimova AA, Notaro A, Arbatsky NP, Speciale I, Shashkov AS, et al. *Acinetobacter baumannii* K13 and K73 capsular polysaccharides differ only in K-unit side branches of novel non-2-ulosonic acids: di-N-acetylated forms of either acinetaminic acid or 8-epiacinetaminic acid. Carbohydr Res. 2017;452:149–55.

53. Kenyon JJ, Arbatsky NP, Sweeney EL, Shashkov AS, Shneider MM, Popova AV, et al. Production of the K16 capsular polysaccharide by *Acinetobacter baumannii* ST25 isolate D4 involves a novel glycosyltransferase encoded in the KL16 gene cluster. Int J Biol Macromol. 2019;128:101–6.

54. Shashkov AS, Kenyon JJ, Arbatsky NP, Shneider MM, Popova AV, Miroshnikov K, et al. Related structures of neutral capsular polysaccharides of *Acinetobacter baumannii* isolates that carry related capsule gene clusters KL43, KL47, and KL88. Carbohydr Res. 2016;435:173–9.

55. Shashkov AS, Arbatsky NP, Senchenkova SN, Kasimova AA, Dmitrenok AS, Shneider MM, et al. Characterization of the carbapenem-resistant *Acinetobacter baumannii* clinical reference isolate BAL062 (CC2: KL58: OCL1): resistance properties and capsular polysaccharide structure. mSystems. 2024;9(10):e00941–24.

56. Shashkov AS, Cahill SM, Arbatsky NP, Westacott A, Kasimova AA, Shneider MM, et al. *Acinetobacter baumannii* K116 capsular polysaccharide structure is a hybrid of the K14 and revised K37 structures. Carbohydr Res. 2019;484:107774.

57. Kasimova AA, Cahill SM, Shpirt AM, Dudnik AG, Shneider MM, Popova AV, et al. The K139 capsular polysaccharide produced by *Acinetobacter baumannii* MAR17-1041 belongs to a group of related structures including K14, K37 and K116. Int J Biol Macromol. 2021;193:2297–303.

58. Shashkov AS, Kenyon JJ, Arbatsky NP, Shneider MM, Popova AV, Miroshnikov K, et al. Structures of three different neutral polysaccharide of *Acinetobacter baumannii*, NIPH190, NIPH201, and NIPH615, assigned to K30, K45, and K48 capsule types, respectively, based on capsule biosynthesis gene clusters. Carbohydr Res. 2015;417:81–8.

59. Shashkov AS, Kenyon JJ, Senchenkova SN, Shneider MM, Popova AV, Arbatsky NP, et al. *Acinetobacter baumannii* K27 and K44 capsular polysaccharides have the same K unit but different structures due to the presence of distinct *wzy* genes in otherwise closely related K gene clusters. Glycobiology. 2016;26(5):501–8.

60. Shpirt AM, Harmer CJ, Shashkov AS, Shneider MM, Chizhov AO, Dmitrenok AS, et al. *Acinetobacter baumannii* nasal carriage isolate recovered from an asymptomatic patient in Vietnam is extensively antibiotic resistant and produces a rare K71 type capsule. Microbiol Spectr. 2024;12(12):e01838–24.

61. Kenyon JJ, Senchenkova SN, Shashkov AS, Shneider MM, Popova AV, Knirel YA, et al. K17 capsular polysaccharide produced by *Acinetobacter baumannii* isolate G7 contains an amide of 2-acetamido-2-deoxy-D-galacturonic acid with D-alanine. Int J Biol Macromol. 2019;144:857–62.

62. Vinogradov E, Zou L, Stupak J, Martynova Y, Arbour M, St Michael F, et al. Capsular Polysaccharide of *Acinetobacter baumannii* MRSN 31196 (a KL1 Variant Strain) and its Degradation by a Recombinant Depolymerase from Bacteriophage vB_AbaP_B5. Carbohydr Res. 2025;556:109621.

63. Mostowy R, Holt K. Diversity-generating machines: Genetics of bacterial sugar-coating. Trends Microbiol. 2018;26(12):1008–21.

64. Wyres K, Wick R, Gorrie C, Jenney A, Follador R, Thomson N, et al. Identification of *Klebsiella* capsule synthesis loci from whole genome data. Microb Genom. 2016;2(12):e000102.

65. Lam MMC, Wick RR, Judd LM, Holt KE, Wyres KL. Kaptive 2.0: updated capsule and LPS locus typing for the *Klebsiella pneumoniae* species complex. Microb Genom. 2022;8(3):000800.

66. Gladstone RA, Pesonen M, Pöntinen AK, Mäklin T, MacAlasdair N, Thorpe H, et al. Identification of transporter-dependent capsular loci associated with the invasive potential of *Escherichia coli*. Nat Microbiol. 2026;11(1205–1216):1–12.

67. Miravet-Verde S, Cacace E, Mores CR, Rutschmann C, Lin CW, Ruscheweyh HJ, et al. *In silico* typing maps the natural diversity of *Escherichia coli* transporter-dependent capsules. Nat Microbiol. 2026;11(5):1217–32.

68. Epping L, Van Tonder AJ, Gladstone RA, Consortium GPS, Bentley SD, Page AJ, et al. SeroBA: rapid high-throughput serotyping of *Streptococcus pneumoniae* from whole genome sequence data. Microb Genom. 2018;4(7):p.e000186.

69. Senchenkova SN, Shashkov AS, Popova AV, Shneider MM, Arbatsky NP, Miroshnikov K, et al. Structure elucidation of the capsular polysaccharide of *Acinetobacter baumannii* AB5075 having the KL25 capsule biosynthesis locus. Carbohydr Res. 2015;408:8–11.

70. Arbatsky NP, Kasimova AA, Shashkov AS, Shneider MM, Popova AV, Hall RM, et al. Carbapenem-resistant *Acinetobacter baumannii* isolate BAL114 from Vietnam produces capsular polysaccharide with two forms of the K58 unit. Carbohydr Res. 2025;556:109592.

71. Arbatsky NP, Shneider MM, Kenyon JJ, Shashkov AS, Popova AV, Miroshnikov KA, et al. Structure of the neutral capsular polysaccharide of *Acinetobacter baumannii* NIPH146 that carries the KL37 capsule gene cluster. Carbohydr Res. 2015;413:12–5.

72. Kenyon JJ, Hall RM, De Castro C. Structural determination of the K14 capsular polysaccharide from an ST25 *Acinetobacter baumannii* isolate, D46. Carbohydr Res. 2015;417:52–6.

73. Kasimova AA, Shneider MM, Arbatsky NP, Popova AV, Shashkov AS, Miroshnikov K, et al. Structure and gene cluster of the K93 capsular polysaccharide of *Acinetobacter baumannii* B11911 containing 5-N-Acetyl-7-N-[(R)-3-hydroxybutanoyl]pseudaminic acid. Biochem (Mosc). 2017;82(4):483–9.

74. Arbatsky NP, Kasimova AA, Shashkov AS, Shneider MM, Popova AV, Shagin D, et al. Structure of the K128 capsular polysaccharide produced by *Acinetobacter baumannii* KZ-1093 from Kazakhstan. Carbohydr Res. 2019.

