## Supplementary Figure 1 and Figure 13 for "Diversity at the *Acinetobacter baumannii* K locus: towards a comprehensive *in silico* database for prediction of capsular polysaccharide types"

Johanna J. Kenyon[1,2,]

[1] *School of Pharmacy and Medical Sciences, Health Group, Griffith University, Parklands Drive, Gold Coast, Queensland, Australia*

[2] *Institute for Biomedicine and Glycomics, Griffith University, Parklands Drive, Gold Coast, Queensland, Australia.*

### Table of Contents

**SUPPLEMENTARY FIGURES**
**Supplementary Figure 1.** Release of *A. baumannii* genomes into National Centre for Biotechnology Innovation (NCBI) databases (March 2007 to January 2026)
**Supplementary Figure 2.** *A. baumannii* K loci that harbour *psa* genes
**Supplementary Figure 3.** *A. baumannii* K loci that harbour *lga* genes
**Supplementary Figure 4.** *A. baumannii* K loci that harbour *neu* genes
**Supplementary Figure 5.** *A. baumannii* K loci that harbour *gna1-gne2* genes
**Supplementary Figure 6.** *A. baumannii* K loci that harbour *dga* genes
**Supplementary Figure 7.** *A. baumannii* K loci that harbour *rmlBDAC* genes
**Supplementary Figure 8.** *A. baumannii* K loci that harbour *fdt* or *qdt* genes
**Supplementary Figure 9.** *A. baumannii* K loci that harbour *rhnABC* genes
**Supplementary Figure 10.** *A. baumannii* K loci that harbour *vio* genes
**Supplementary Figure 11.** *A. baumannii* K loci that harbour *mna* genes
**Supplementary Figure 12.** *A. baumannii* K loci that harbour *gna-wzx* gene modules
**Supplementary Figure 13.** Groups of K loci that differ in Region 3 gene content.

**SUPPLEMENTARY TABLES**
**Supplementary Table 1.** *A. baumannii* genome assemblies used in this study.
**Supplementary Table 2.** *A. baumannii* KL reference sequences included in the database.
**Supplementary Table 3.** Details of annotations ascribed to CPS biosynthesis genes detected in *A. baumannii*
**Supplementary Table 4.** *Kaptive* output for 46,185 genomes screened against custom database with 241 KL
**Supplementary Table 5.** Summary of gene presence/absence matrix generated for 409 KL using *PPanGGOLiN* with an 85% translated identity threshold.
**Supplementary Table 6.** *Kaptive* output for 46,185 genomes screened against database with 409 KL and 10 extra-locus genes

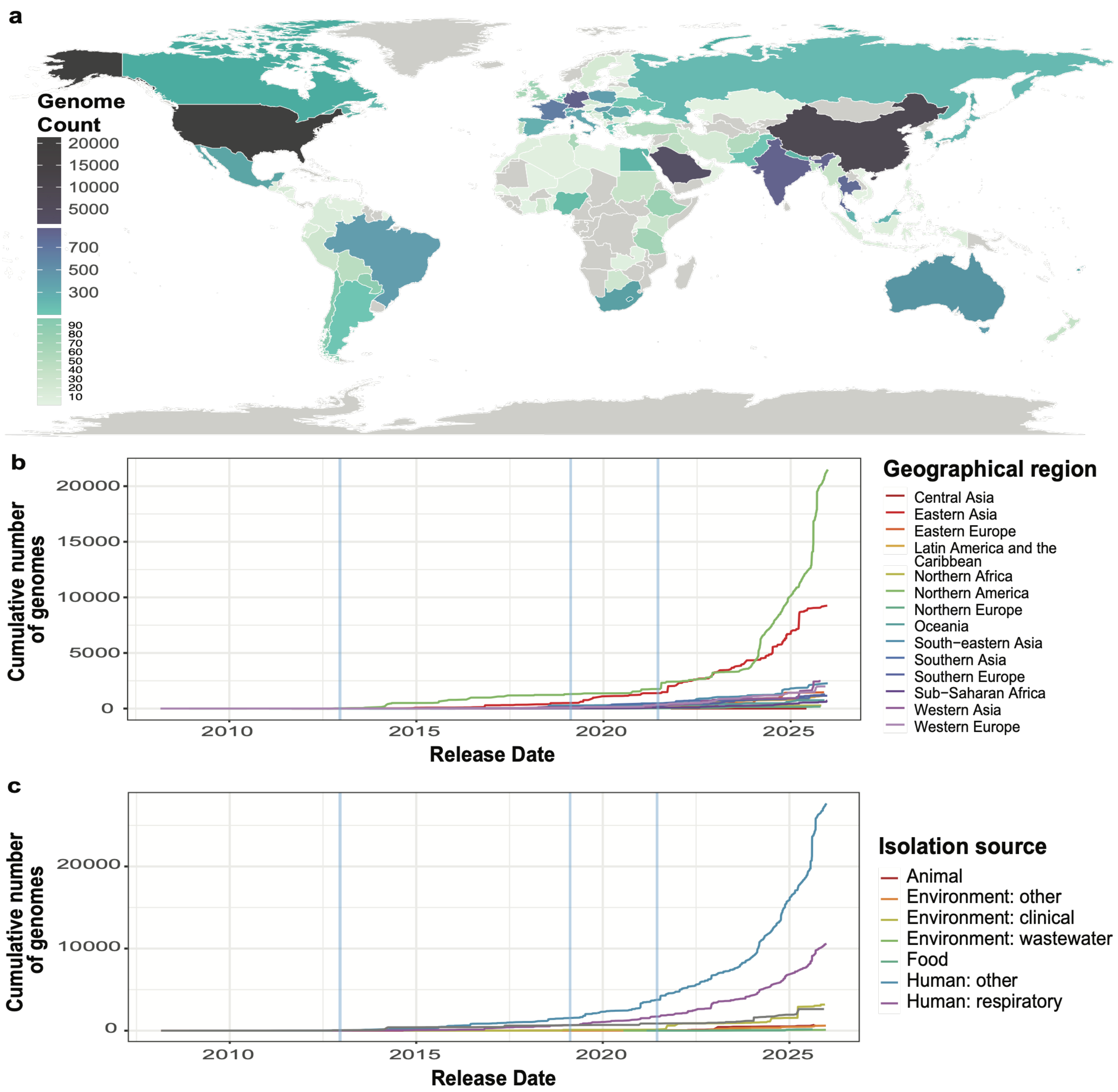


**Supplementary Figure 1**. Release of *A. baumannii* genomes into National Centre for Biotechnology Innovation (NCBI) databases (March 2007 to January 2026), indicating the proportion of total genomes per **(a)** country, and cumulative number of genomes per **(b)** UNESCO region if known, and **(c)** source per year if known. Blue vertical bars in bar charts indicate key studies reporting characterisation and annotation of *A. baumannii* K loci conducted in 2013 (19), 2019 (18) and 2021 (17).

| | None | *gne1* | *gne1/ pgt1* | *pgt1* | *gne1/ pet1* | *gne1/ atr12* | *gne1/ atr15* | *gne1/ atr20* | *gne1/ orf/ atr20** | *gne1/ orf/ atr36* | *gne1/ atr42/ atr43* | *gne1/ atr58/ pgt1* | *gne1/ atr60/ pgt1* |
|---|---|---|---|---|---|---|---|---|---|---|---|---|---|
| 1 | KL1 | KL107 | | | | | | | | | | | |
| 2 | | KL2 | KL81 | | | | | | | | | | |
| 3 | | KL3 | KL22 | | | | | | KL159 | | | | |
| 4 | KL109 | KL9 | KL149 | | | | | KL168 | KL173 | | | | |
| 5 | | KL290 | KL10 | | | | | | | | | | |
| 6 | | | KL195 | KL11 | | | | | | | | | |
| 7 | | KL309 | KL14 | | | | | | | | | | |
| 8 | KL147 | KL15 | | | | | | | | | | | |
| 9 | | KL16 | | | | KL351 | | | | | | | |
| 10 | KL17 | KL18 | KL237 | | | | | | | | | | |
| 11 | | KL34 | KL199 | | KL20 | | | | | | | | |
| 12 | KL201 | KL25 | | | | | | | | | | | |
| 13 | | KL304 | KL27 | | | KL130 | | | | | | | |
| 14 | | KL32 | KL200 | | | | KL100 | KL164 | | | | | |
| 15 | | KL33 | KL357 | | | | | KL77 | | | | | |
| 16 | | | KL345 | KL36 | | | | | | | | | |
| 17 | | KL346 | KL37 | | | | | | | | | | |
| 18 | KL91 | KL40 | KL250 | | | | | | | | | | |
| 19 | | KL259 | | | | KL41 | | | | | | | |
| 20 | | KL42 | KL347 | | | | | | | | KL216 | | |
| 21 | | KL249 | KL44 | | | | | | | | | | |
| 22 | | KL231 | KL47 | | | | | | | | | | |
| 23 | | KL150 | | | | | KL50 | | | | | | |
| 24 | | KL261 | KL51 | | | | | | | | | | |
| 25 | | KL196 | KL52 | | | | | | | | | | |
| 26 | KL53 | KL350 | | | | | | | | | | | |
| 27 | | KL404 | | | | | | | | | KL58 | | |
| 28 | | KL64 | KL160 | | | | | | | | | | |
| 29 | | | | | | | | KL124 | KL82 | | | | |
| 30 | | | KL359 | KL83 | | | | | | | | | |
| 31 | | KL101 | KL295 | | | | | | | | | | |
| 32 | | KL296 | KL104 | | | | | | | | | | |
| 33 | | KL161 | | | KL118 | | | | | | | | |
| 34 | | KL138 | KL122 | | | | | | | | | | |
| 35 | KL299 | KL126 | | | | | | | | | | | |
| 36 | | | KL129 | KL220 | | | | | | | | | |
| 37 | | KL152 | | | | | | | KL151 | | | | |
| 38 | | KL155 | KL210 | | | | | | | | | | |
| 39 | | KL170 | KL225 | | | | | | | | | | |
| 40 | | | KL198 | KL269 | | | | | | | | | |
| 41 | KL202 | KL322 | | | | | | | | | | | |
| 42 | | KL226 | KL273 | | | | | | | | | | |
| 43 | | KL233 | KL334 | | | | | | | | | | |
| 44 | | KL238 | KL260 | | | | | | | | | | |
| 45 | | KL254 | | | | | | | | KL122 | | | |
| 46 | | | KL335 | KL244 | | | | | | | | | |
| 47 | | KL256 | | | | | | | | | | KL340 | KL384 |
| 48 | | KL314 | KL262 | | | | | | | | | | |
| 49 | KL307 | KL366 | | | | | | | | | | | |
| 50 | KL301 | KL403 | | | | | | | | | | | |
| 51 | | KL342 | KL349 | | | | | | | | | | |
| 52 | KL329 | KL328 | | | | | | | | | | | |
| 53 | | KL333 | | | KL401 | | | | | | | | |

**Supplementary Figure 13. Groups of K loci that differ in Region 3 gene content.** Heatmap showing variation of gene content in Region 3 for 53 groups found. Gene combinations are shown above, and those with an elucidated CPS structure are highlighted in blue.
