## Supplementary figures and images for "Diversity at the *Acinetobacter baumannii* K locus: towards a comprehensive *in silico* database for prediction of capsular polysaccharide types"

### FigS2_psa.pdf

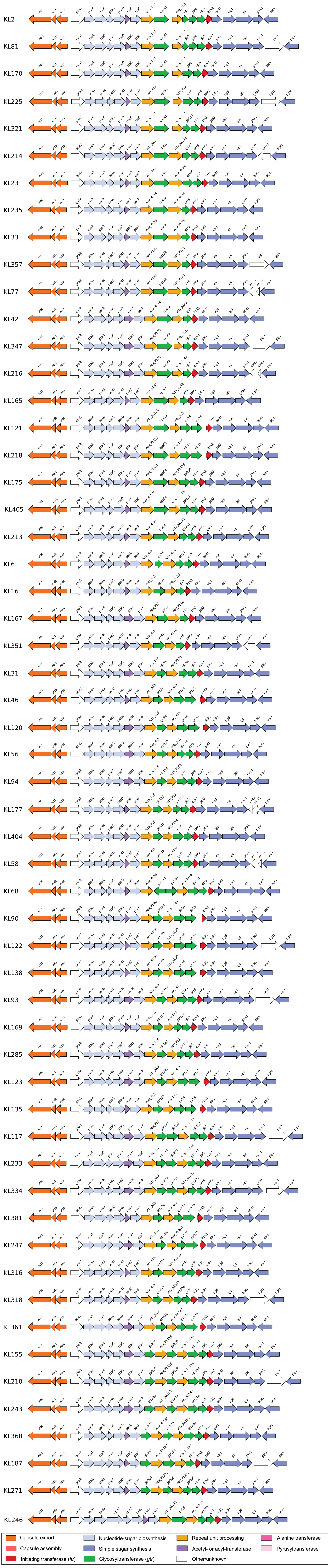

### FigS3_lga.pdf

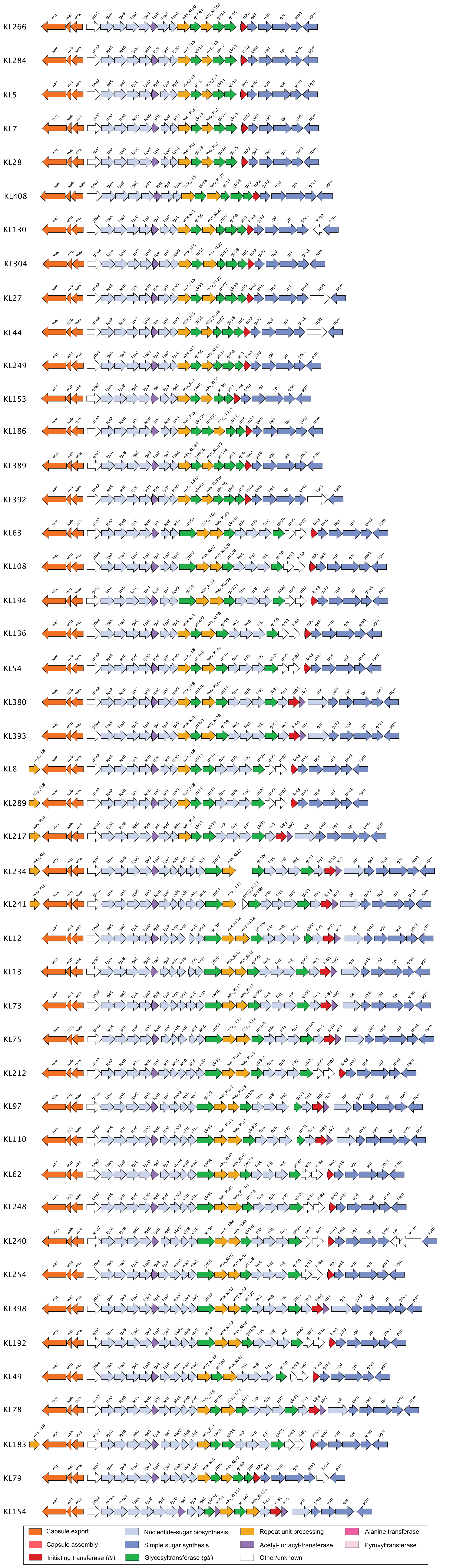

### FigS4_neu.pdf

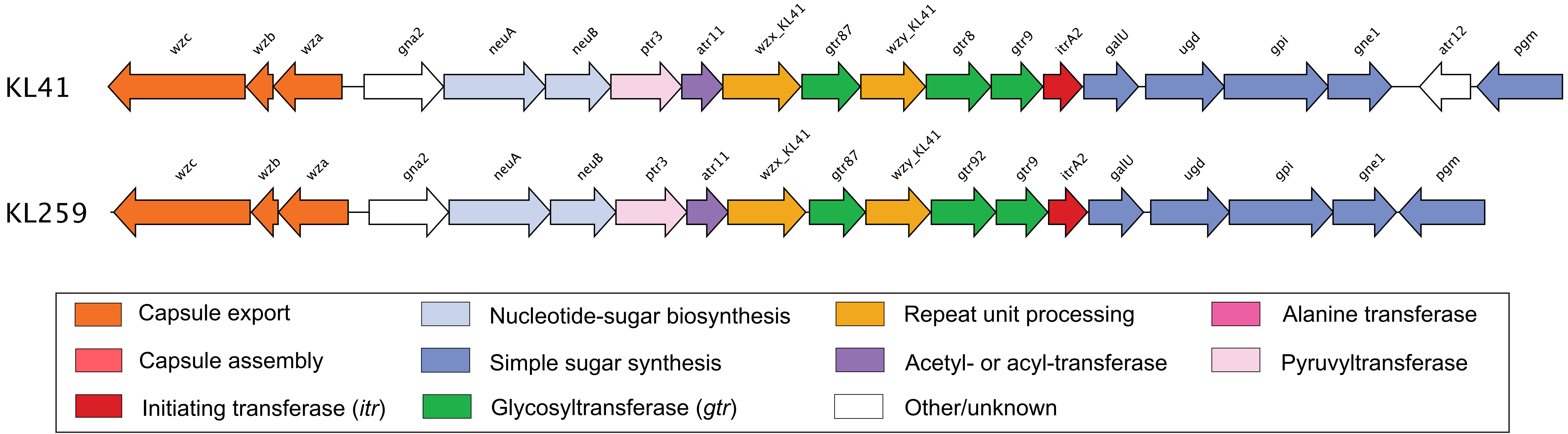

### FigS5_gna_gne2.pdf

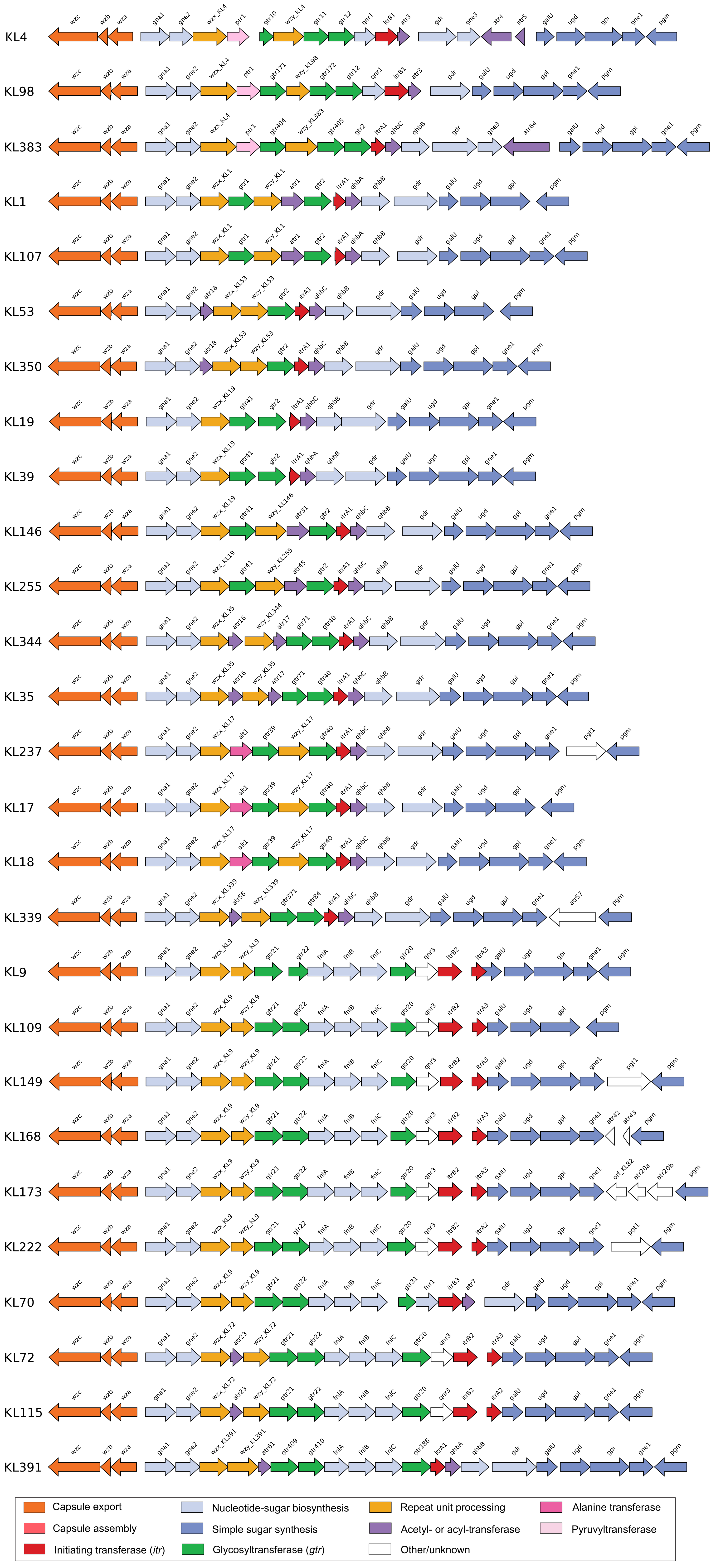

### FigS6_dga.pdf

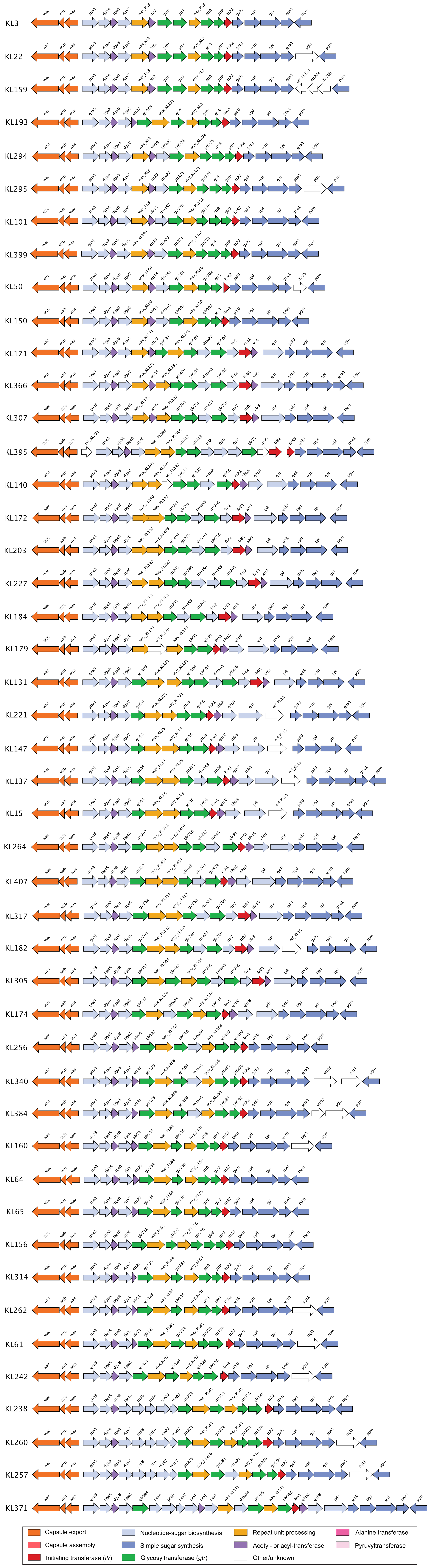

### FigS7_rmlBDAC.pdf

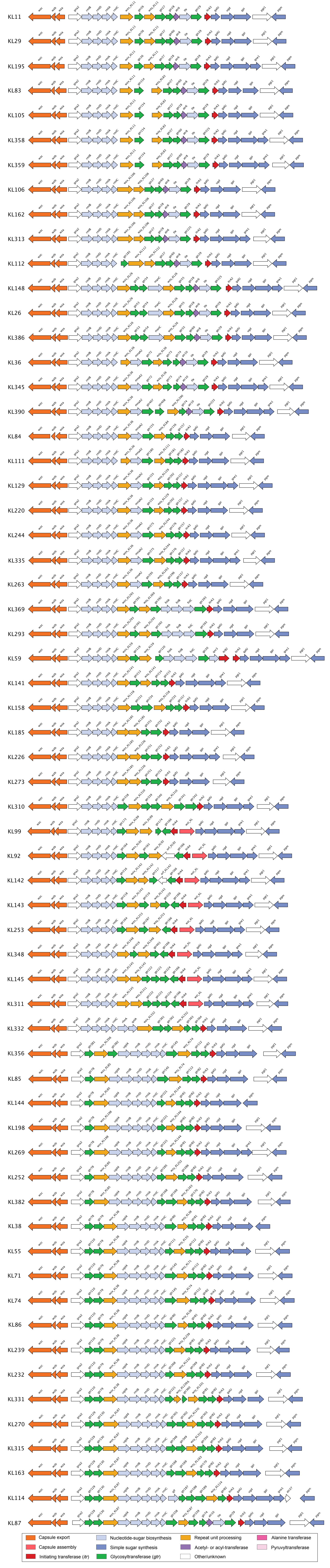

### FigS8_fdt-qdt.pdf

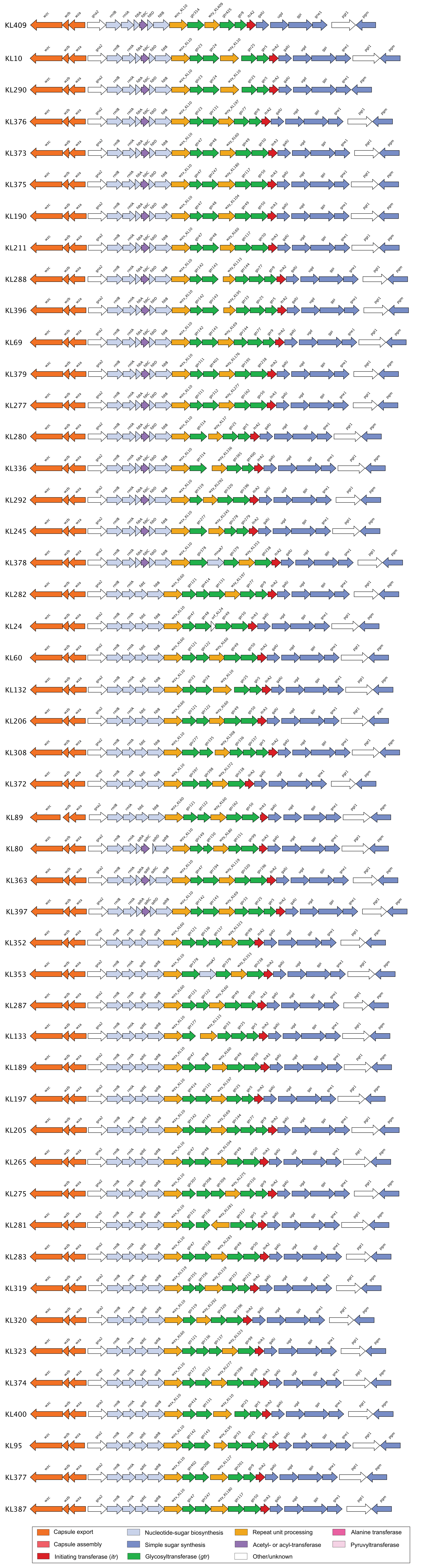

### FigS9_rhnABC.pdf

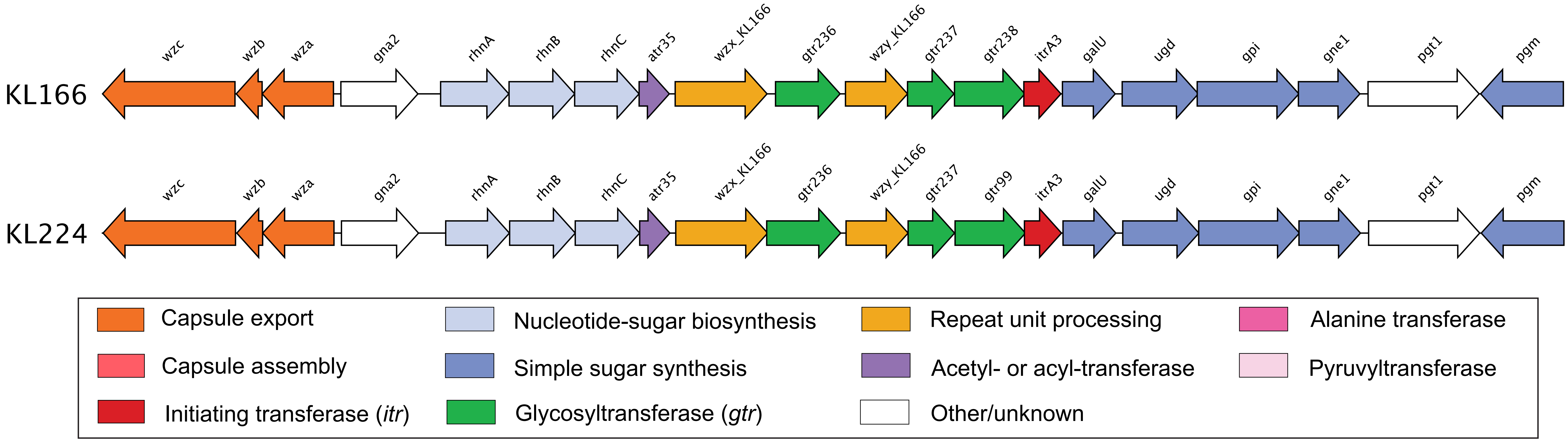

### FigS10_vio.pdf

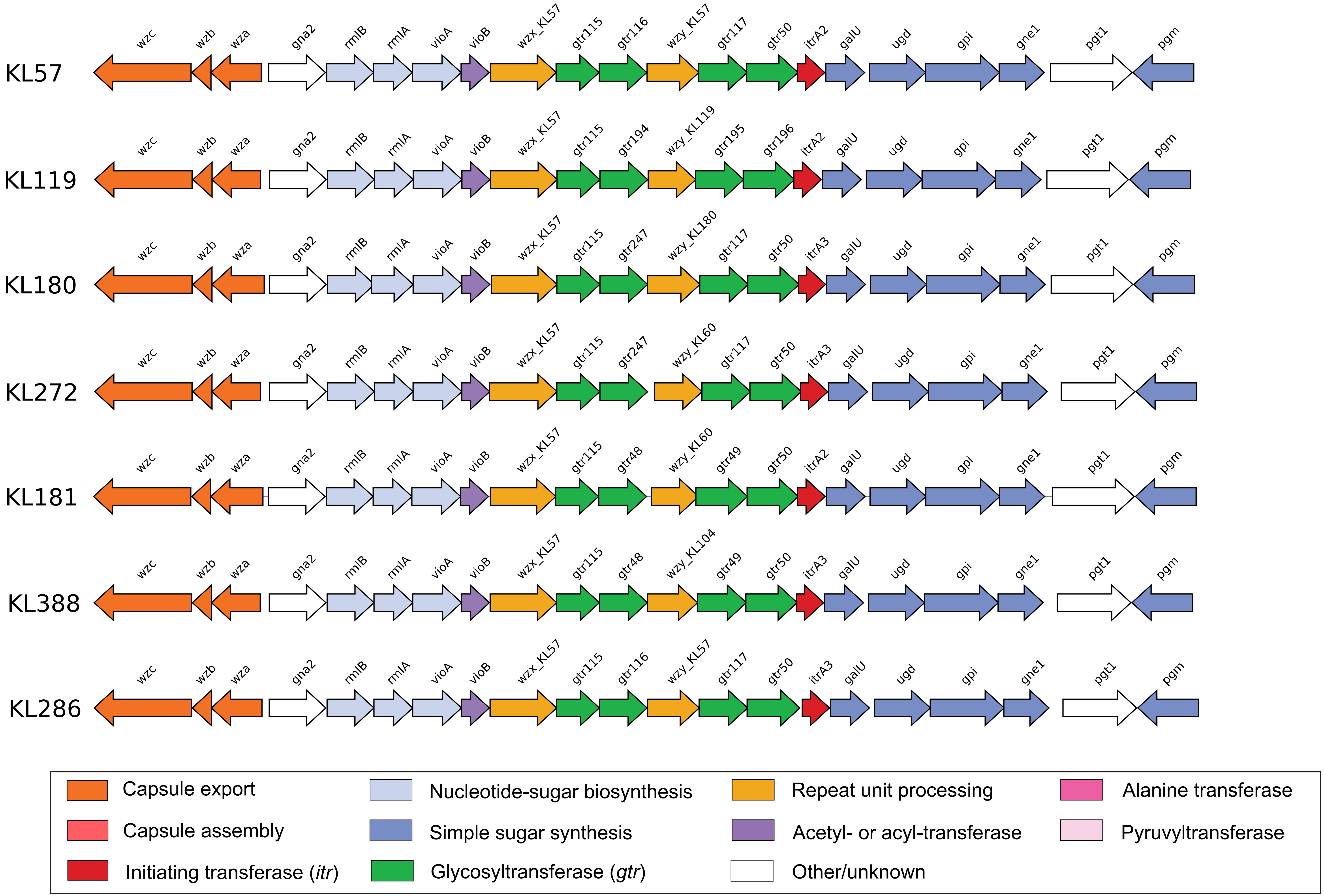

### FigS11_mna.pdf

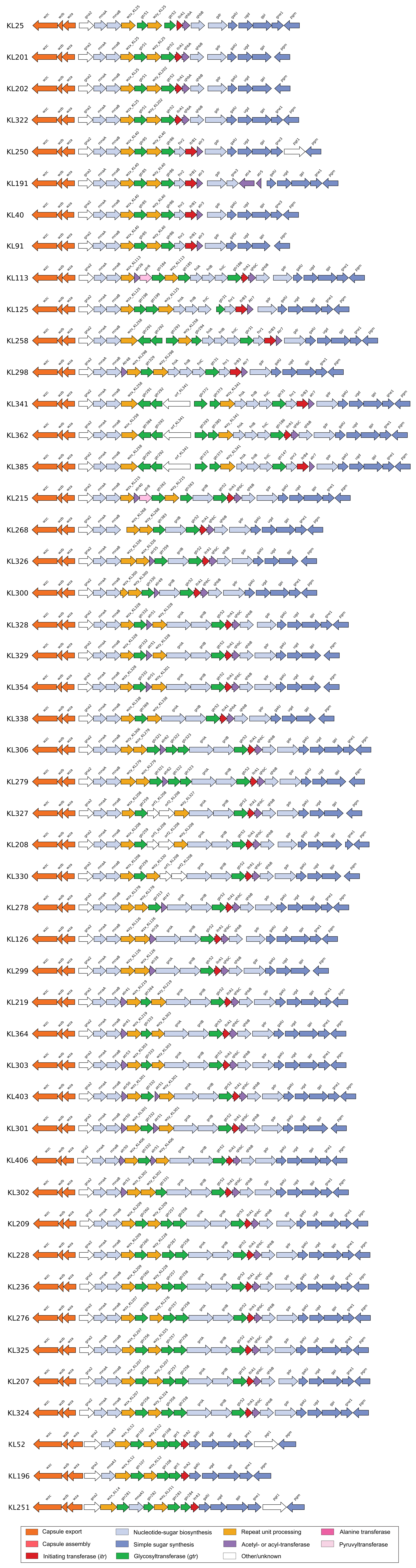

### FigS12_gna-wzx.pdf

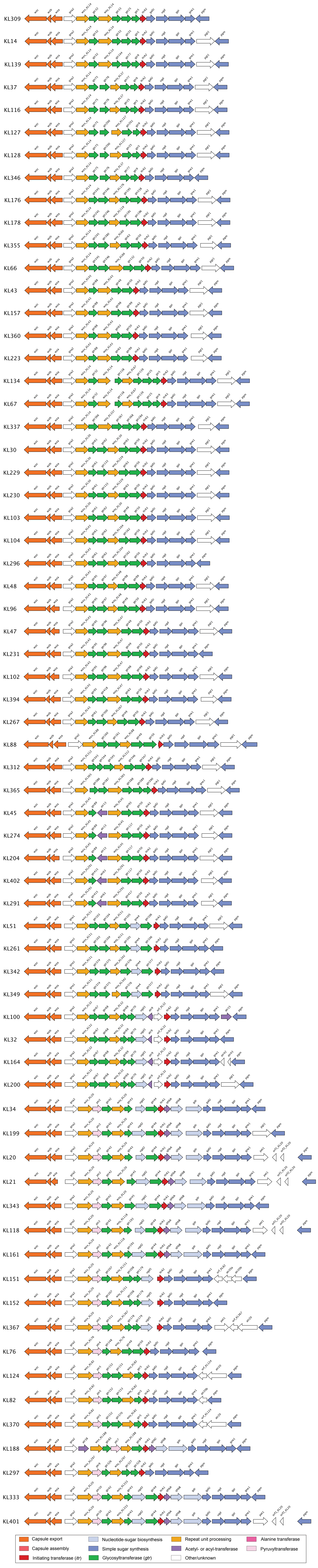
